# Identification of a somatic H3K23me3 methyltransferase SET-19 in *C. elegans*

**DOI:** 10.64898/2026.03.24.713969

**Authors:** Mingjing Xu, Zixue Fan, Chaoyue Yan, Xiangyang Chen, Xinya Huang, Chengming Zhu, Minjie Hong, Jiewei Cheng, Xinhao Hou, Shuju Li, Mengfeng Li, Zhongying Zhao, Yunyu Shi, Meng Huang, Shouhong Guang, Xuezhu Feng

## Abstract

Histone methylation plays essential roles in modulating chromatin organization and gene expression. H3K23 methylation is a conserved histone modification, yet its biological roles and the enzymes responsible for its deposition remain poorly understood. Here, we show that the loss of *set-19* leads to a pronounced reduction in H3K23 methylation in *C. elegans*, as revealed by quantitative mass spectrometry, western blotting, and immunofluorescence staining. In vitro biochemical assays show that recombinant SET-19 proteins purified from *E. coli* directly catalyze H3K23 methylation. Genome-wide chromatin immunoprecipitation assays reveal that H3K23me3 is enriched at heterochromatic regions and that loss of *set-19* alters H3K23me3 levels, accompanied by derepression of a subset of genes. Genetic analyses indicate that SET-19 is dispensable for both germline and somatic RNAi as well as transgenerational epigenetic inheritance of RNAi. SET-19 is predominantly expressed in somatic cells and specifically mediates H3K23me3 deposition in somatic tissues. The loss of *set-19* causes a modest developmental delay. Together, our results identify SET-19 as a somatic H3K23 methyltransferase and link SET-19-dependent H3K23me3 to transcriptional repression in *C. elegans*.

## Introduction

The nucleosome, composed of DNA wrapped around histone proteins (H2A, H2B, H3, and H4), is the fundamental structural unit of chromatin. Post-translational modifications (PTMs), such as methylation, acetylation, and phosphorylation, occur on both histone tails and core regions ^1^. Histone modifications function as epigenetic marks that regulate chromatin organization and stability, as well as DNA replication, repair, and transcription.

Chromatin is broadly divided into euchromatin and heterochromatin, which differ in their degrees of compaction and transcriptional activity. Histone PTMs such as acetylation, H3K4 methylation, and H3K36 methylation are typically enriched in transcriptionally active euchromatic regions and are associated with gene activation. In contrast, repressive marks including H3K9 and H3K27 methylation are preferentially enriched in transcriptionally silent heterochromatic regions. The distinct genomic distributions of these modifications reflect their diverse regulatory functions, and the dynamic regulation of these marks across tissues and developmental stages contributes to cell fate determination, fertility, development, and epigenetic inheritance ^2-6^.

Histone PTMs are dynamically regulated by specific writers, readers, and erasers, which establish, recognize, and remove these modifications, respectively ^1,7^. Writers responsible for histone methylation are histone methyltransferases (HMTs), which catalyze the methylation of lysine (K) and arginine (R) residues on histone proteins. Lysine residues can be mono-, di-, or trimethylated by lysine methyltransferases (KMTs), whereas arginine residues can be mono- or dimethylated by arginine methyltransferases (RMTs) ^8^. Most KMTs harbor a conserved SET domain that confers catalytic activity and substrate specificity ^7,9-11^.

Histone H3 lysine 23 methylation (H3K23me) is widely conserved across eukaryotic species ^12-24^. In organisms including *Tetrahymena*, *C. elegans*, and mammals, evidence from immunostaining, chromatin immunoprecipitation (ChIP)-based analyses, and binding studies with HP1 family proteins supports an association of H3K23 methylation with heterochromatic regions ^14,15,17,18,20,21,24-26^. In *Tetrahymena*, the loss of the H3K23 methyltransferase Ezl3p leads to aberrant targeting of meiosis-induced DNA double-strand breaks to heterochromatin and compromises progeny viability ^17^. In *C. elegans*, H3K23me3 shares genomic features with classical repressive marks such as H3K9 and H3K27 methylation and is also associated with transgenerational epigenetic silencing and accumulation at RNAi target loci ^25-30^. H3K9 methylation, deposited primarily by the SET-domain proteins MET-2 and SET-25, contributes to satellite repeat repression and the maintenance of tissue-specific gene expression patterns ^3,31^. H3K27 methylation, mediated by MES-2, is also involved in RNAi-associated chromatin silencing ^30^. The loss of the corresponding histone methyltransferases for H3K9me3 and H3K27me3 results in varying degrees of RNAi defects ^30,32-34^. Two methyltransferases, SET-32 and SET-21, have been reported to contribute to H3K23 methylation ^25,26^. However, loss of either enzyme alone has not been shown to cause a substantial global reduction in H3K23 methylation in vivo, which may reflect their tissue- or stage-specific contributions and potential functional redundancy among these enzymes or with other H3K23 methyltransferases ^25,26,35^. Despite these advances, the enzymatic basis, tissue-specific regulation, and biological functions of H3K23 methylation remain far less well characterized than those of other histone methylation marks. The *C. elegans* genome encodes 38 SET domain-containing proteins ^36-47^, many of which remain functionally uncharacterized. Our previous work showed that the loss of *set-19* leads to a significant reduction in H3K23me3 levels in vivo ^48^, implicating SET-19 as a candidate H3K23 methyltransferase.

In this work, we investigated the role of SET-19 in H3K23 methylation in *C. elegans* and showed that SET-19 functions as a somatic H3K23 methyltransferase required for H3K23me3 methylation in vitro and in vivo. This work provides insights into how H3K23 methylation is established and supports the idea that distinct methyltransferases may function in different tissues or cell types to promote proper development.

## Results

### SET-19 is required for H3K23 methylation in *C. elegans*

In *C. elegans,* the biological functions of many SET domain-containing methyltransferases remain poorly understood ^25,26,36-46^. SET-19 harbors a conserved SET domain (Fig. 1a). Our previous work showed that the *set-19(ok1813)* mutant exhibited reduced levels of H3K23me2 and H3K23me3, as detected by western blotting^48^. However, the role of SET-19 in H3K23 methylation required further validation.

**Fig. 1.**
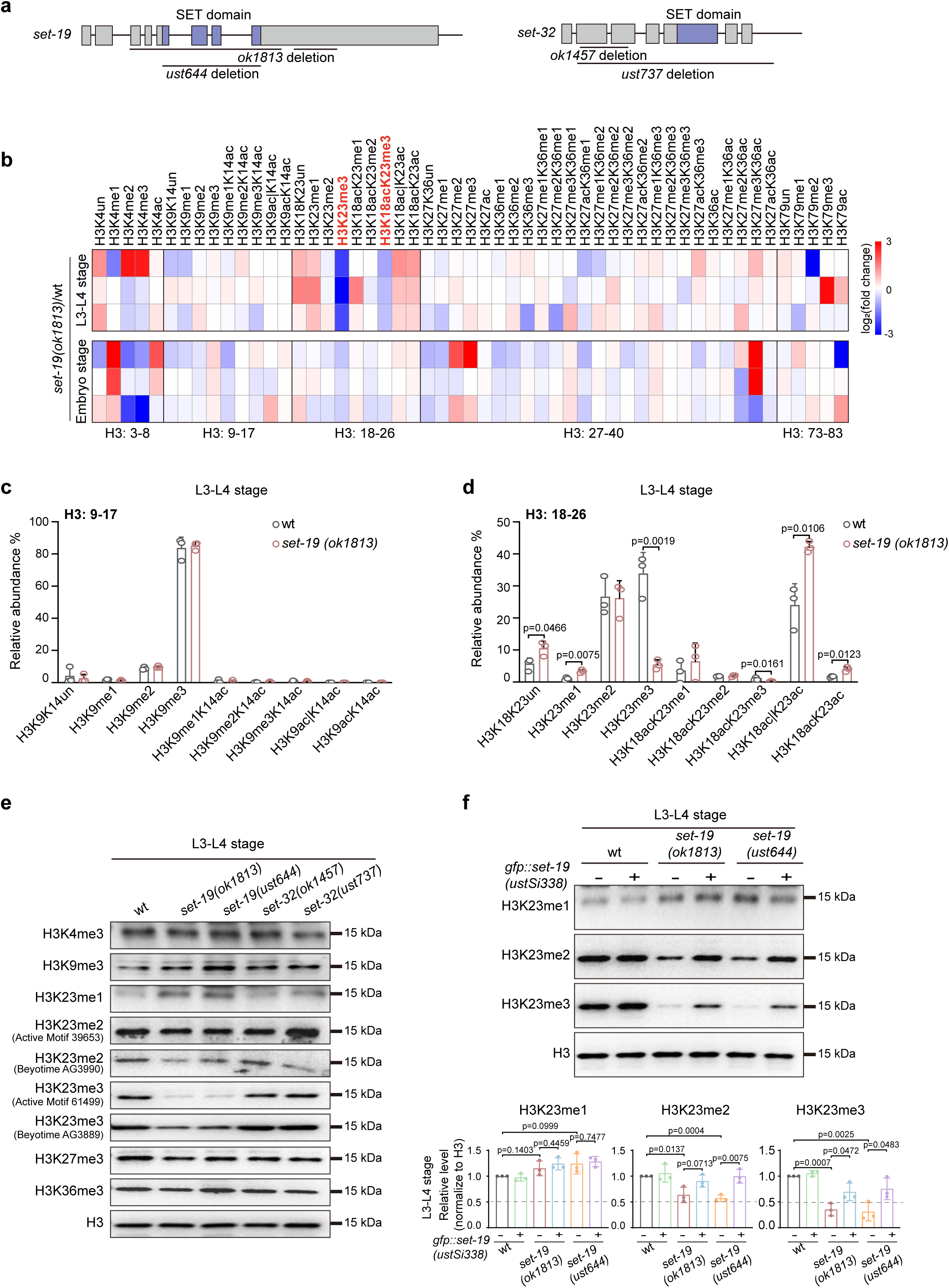
Loss of SET-19 leads to a marked reduction of H3K23me3. **a,** Schematic representation of the *set-19* and *set-32* genes. The *set-19(ok1813)* allele contains two large deletions and likely functions as a null allele. The *set-19(ust644)* allele carries a precise deletion of the SET domain, and the open reading frame is retained. The *set-32(ok1457)* allele contains a deletion that does not disrupt the open reading frame (ORF), whereas *set-32(ust737)* harbors a large deletion and likely functions as a null allele. SET domains are indicated in purple. **b,** Heatmap showing relative abundances of histone H3 post-translational modifications (PTMs) measured by mass spectrometry. Values represent log₂(fold change) in *set-19(ok1813)* mutants relative to wild-type (wt) animals for each biological replicate. **c, d,** Quantification of H3(9–17) and H3(18–26) methylation states in wild-type and *set-19(ok1813)* animals at the indicated developmental stages. Methylation levels are shown as relative abundances, calculated by dividing the AUC of each modified peptide by the sum of the AUCs of all detected forms of that peptide and multiplying the resulting ratio by 100. Statistical significance was assessed using a two-tailed t test, with P < 0.05 considered statistically significant. P values are indicated only for statistically significant differences. *N* = 3 biological replicates. **e,** Western blotting analysis of global H3 methylation levels in L3–L4-stage worms of the indicated genotypes. **f,** Western blotting analysis of H3K23me1/2/3 levels in L3–L4 worms. The *fib-1p::gfp::set-19(ustSi338)* transgene was used for rescue. Signals were normalized to total H3 and expressed relative to wild type. Data are presented as the mean ± SD; N = 3 biological replicates; two-tailed t test. p values are indicated.

To assess the impact of *set-19* loss on histone H3 methylation, we quantified global H3 methylation levels in the *set-19(ok1813)* mutant by mass spectrometry at the L3–L4 larval and embryonic stages. The H3K23me3 levels decreased from approximately 33.8% to 5.4% at the larval stage and from 10% to 8% at the embryonic stage (Fig. 1b–d and Fig. S1f). Mass spectrometry revealed no significant change in H3K23me2, whereas H3K23me1 levels were slightly increased at the L3–L4 larval stage (Fig. 1d). In addition, modest changes in other histone modifications were observed in embryos, including reduced H3K9me2, H3K9me1K14ac, and H3K18acK23me2 levels (p = 0.0386, 0.0161, and 0.0490, respectively), increased H3K9me3 (p = 0.0227), and altered H3K27/K36 combinatorial methylation states (p = 0.0494, 0.0032, and 0.0194; Fig. S1).

To further confirm the role of SET-19, we generated an independent *set-19* allele using CRISPR/Cas9-mediated genome editing. This allele, *set-19(ust644)*, precisely deletes the SET domain without disrupting the open reading frame (ORF) (Fig. 1a). Western blotting analysis revealed that both *set-19(ok1813)* and *set-19(ust644)* mutants exhibited a pronounced reduction in H3K23me3 and a detectable decrease in H3K23me2 at the larval stage (Fig. 1e and Fig. S2). Two independent antibodies were used to detect each of H3K23me2 and H3K23me3 in the western blotting assays. Importantly, reintroduction of a GFP::SET-19 transgene (*fib-1p::gfp::set-19*, *ustSi338*) into the *set-19* mutant background restored H3K23me2 and H3K23me3 levels (Fig. 1f and Fig. S3k). Quantification of western blotting did not reveal significant reductions in H3K23me1, H3K23me2, or H3K23me3 levels in *set-32* single mutant at the L3–L4 larval or embryonic stages (Fig. 1e and Figs. S2 and S3). Nevertheless, these findings do not exclude tissue- or context-specific contributions from SET-32, which may be masked in whole-animal measurements and by the activities of other H3K23 methyltransferases.

Together, these results indicate that SET-19 is required for H3K23 methylation, particularly H3K23me3 in *C. elegans*.

### Recombinant SET-19 catalyzes histone H3K23 methylation in vitro

SET-19 contains a central SET domain and a nearby coiled-coil region, with intrinsically disordered regions (IDRs) at both the N- and C-termini (Fig. 2a). To test whether SET-19 is a bona fide methyltransferase, we performed in vitro histone methylation assays. We attempted to express full-length SET-19 and several SET-19 fragments as GST-fusion proteins in *E. coli*. Although full-length SET-19 could not be successfully expressed and was therefore excluded from subsequent analyses, we obtained purified GST-fused proteins corresponding to the SET domain alone (SET-19 SET; amino acids 37–318) and a fragment containing both the SET and coiled-coil domains (SET-19 SET+CC; amino acids 37–454) (Fig. 2a). SET-32 has previously been reported to exhibit H3K23 methyltransferase activity in vitro and was therefore included as a positive control ^25^. Given the high evolutionary conservation of histone H3 and the conserved nature of lysine 23 across species (Fig. S4a), human histone H3 (Active Motif, #31294) was used as the methylation substrate. Purified GST-SET-19 fusion proteins or full-length GST-SET-32 were incubated with unmodified H3 in the presence of the methyl donor S-adenosylmethionine (SAM). The reaction products were analyzed by western blotting using a panel of antibodies recognizing distinct H3 methylation marks.

**Fig. 2.**
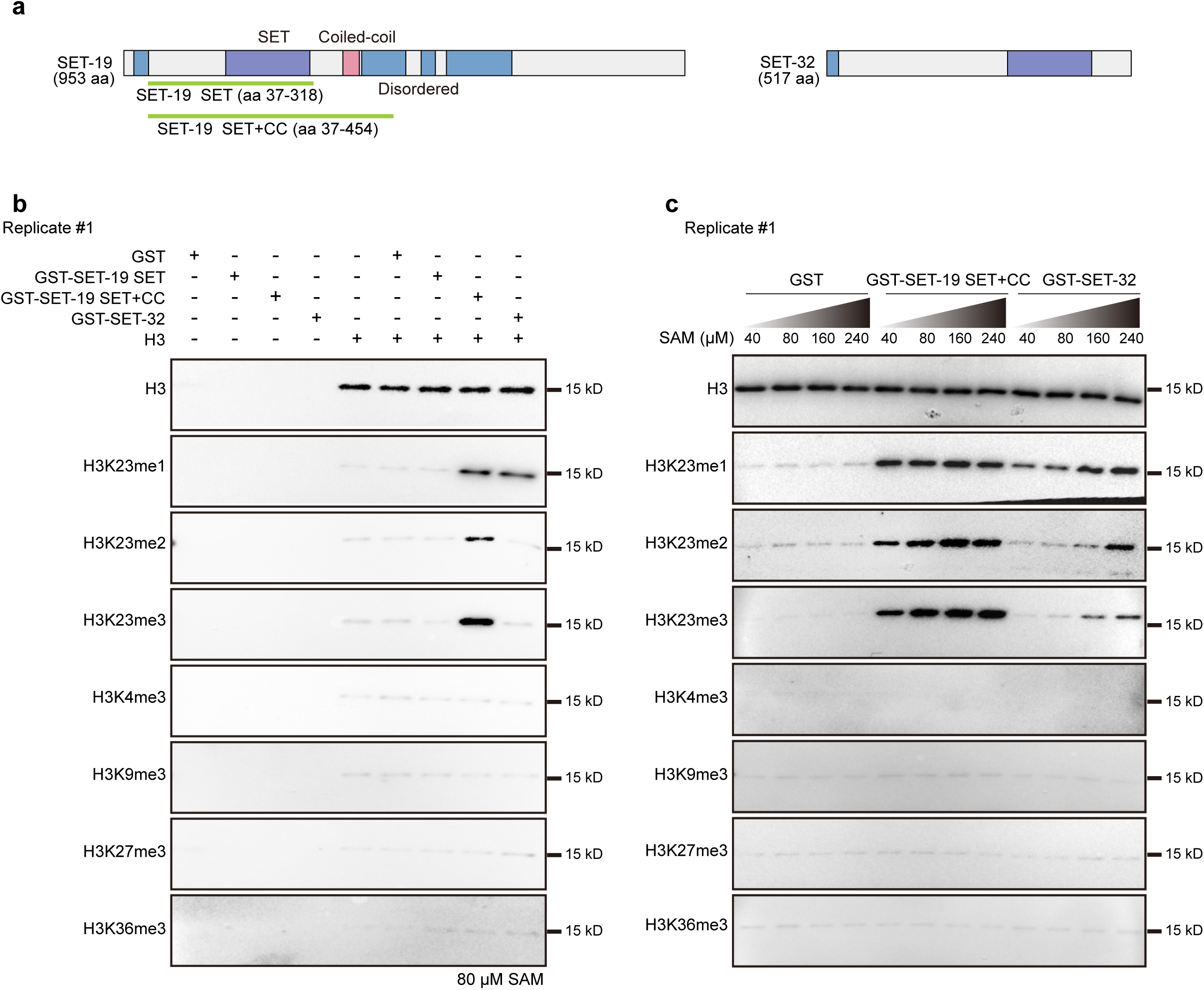
SET-19 specifically catalyzes histone H3K23 methylation in vitro. **a,** Schematic diagram of SET-19 and SET-32 proteins. Purple: SET domain, blue: disordered regions (IDRs), pink: coiled-coil region. Recombinant SET-19 SET or SET+CC fragments are shown in green. **b,** In vitro histone methyltransferase assays using GST-fused SET-19 SET or SET+CC fragments or full-length SET-32 incubated with unmodified human histone H3 and 80 μM S-adenosylmethionine (SAM). Methylation was detected by western blotting. **c,** In vitro methylation assays performed as in (**b**) with increasing concentrations of SAM.

In the in vitro methylation assays, neither the SET-19 fragments nor full-length SET-32 induced detectable changes in H3K4me3, H3K9me3, H3K27me3, or H3K36me3 levels (Fig. 2b and Fig. S4b). In contrast, the SET-19 SET+CC fragment, but not the SET domain alone, induced a pronounced increase in H3K23me1, H3K23me2, and H3K23me3 signals (Fig. 2b and Fig. S4b), indicating that SET-19 specifically methylates H3K23 in vitro and that this activity requires sequences outside the SET domain, which may be important for proper folding or catalytic activity of the SET domain. SET-32 exhibited modest H3K23me1 methyltransferase activity in an in vitro biochemical assay with purified recombinant proteins (Fig. 2b and Fig. S4b), which is consistent with previous reports. In addition, a trend toward reduced H3K23me1/2 levels has been reported in *set-32(smb-11)* mutants ^32^.

SET domain-containing methyltransferases differ not only in substrate specificity but also in their sensitivity to SAM concentration ^41,49-52^. We then examined the enzymatic activities of GST-SET-19 SET+CC and full-length GST-SET-32 across a range of SAM concentrations. Increasing SAM levels enhanced H3K23me1/2/3 signals for both enzymes (Fig. 2c and Fig. S4c). No detectable increases in the tested H3K4me3, H3K9me3, H3K27me3, or H3K36me3 signals were observed under these conditions (Fig. 2c and Fig. S4c). These results indicate that both SET-19 and SET-32 are capable of catalyzing H3K23 methylation in vitro under the tested conditions.

Notably, although SET-19 catalyzed mono-, di-, and trimethylation of H3K23 in vitro, both mass spectrometry and western blotting analyses revealed a predominant reduction in H3K23me3 upon loss of *set-19*. Together, these in vitro and in vivo data establish SET-19 as an H3K23 methyltransferase in *C. elegans* and indicate that SET-19 plays a prominent role in H3K23 trimethylation. SET-32 may also contribute to H3K23 methylation in germline cells or at specific developmental stages, potentially through functional overlap with other H3K23 methyltransferases ^25,26,32,35^.

### SET-19 contributes to genomic H3K23me3 occupancy and the repression of a subset of genes

Previous studies using ChIP-seq and immunofluorescence staining have shown a strong association between H3K23 methylation (H3K23me2/3) and heterochromatin marks ^17,18,20,21,24-26^. In *C. elegans*, the genome-wide distribution of H3K23me3 closely resembles those of H3K9me3 and H3K27me3 (Fig. S5a). *C. elegans* chromosomes are organized into broad domains, with active marks (H3K4me3 and H3K36me3) enriched in central regions and repressive marks (H3K9me1/2/3) enriched in distal arms ^36,53-55^. Consistent with this chromosomal organization, H3K23me3-enriched regions were predominantly localized to chromosome arms (Fig. S5b). In addition, regions centered on H3K23me3 peaks show stronger enrichment of H3K9me3 and H3K27me3 than of the active marks H3K4me3 and H3K36me3 (Fig. 3a).

**Fig. 3.**
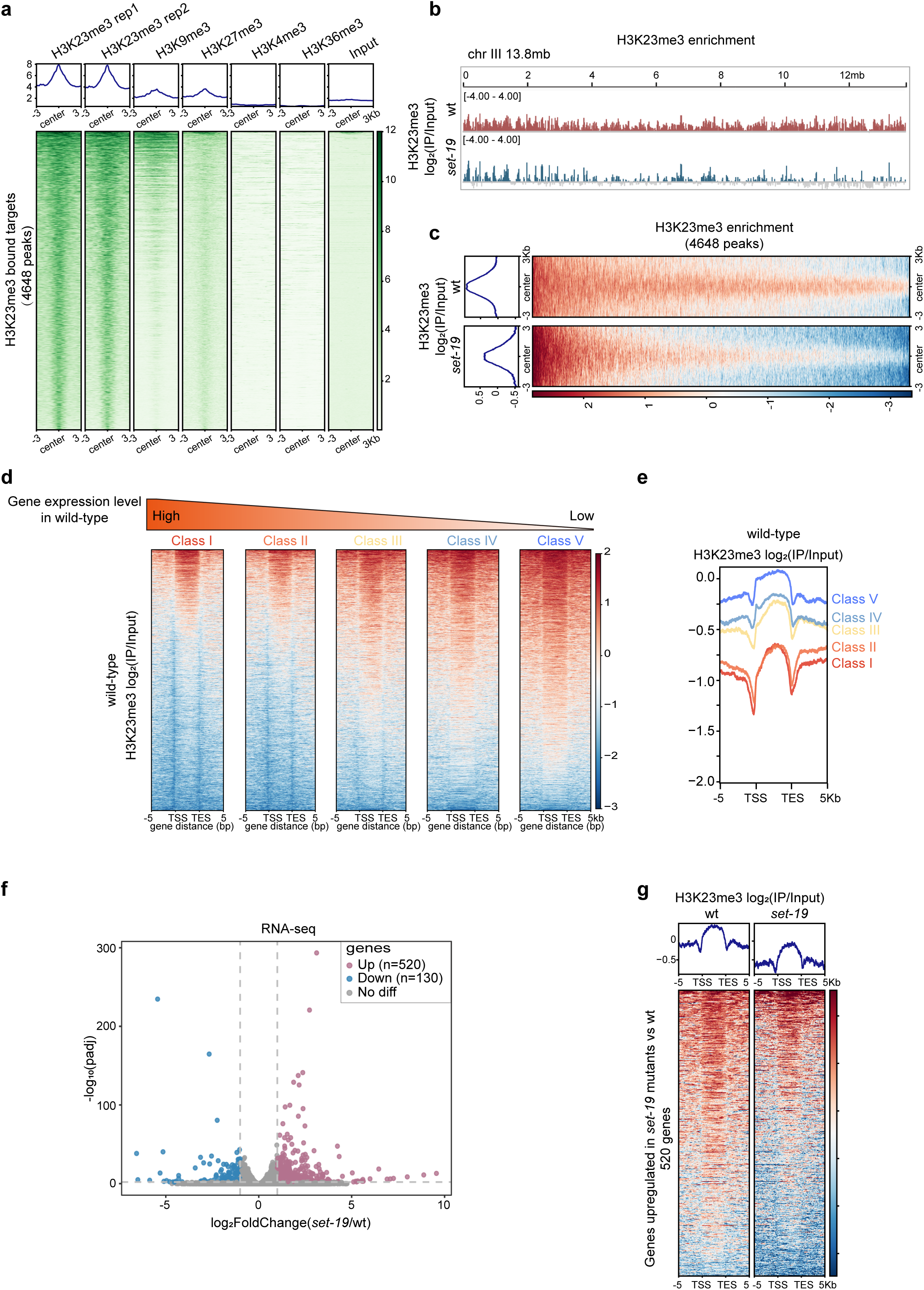
SET-19 contributes to genomic H3K23me3 occupancy and repression of a subset of genes. **a,** Heatmaps showing ChIP-seq signal enrichment of heterochromatic marks (H3K9me3, H3K27me3) and euchromatic marks (H3K4me3, H3K36me3) centered on H3K23me3 peak centers. The average signal within ±3 kb regions is shown. **b,** Genome browser view of H3K23me3 ChIP-seq signals across chromosome III in wild-type and *set-19(ok1813)* animals. Mean log₂ enrichment over input is shown (*N* = 2 biological replicates). H3K23me3 ChIP-seq signals are displayed as log₂(IP/Input) for wild-type and *set-19* samples. **c,** Heatmaps comparing H3K23me3 profiles on H3K23me3 target regions in wild-type and *set-19(ok1813)* worms. Mean log₂ enrichment over input within ±3 kb of peak centers is shown. **d,** Heatmaps of H3K23me3 enrichment across gene bodies in wild-type worms. Genes were grouped into quintiles based on RNA-seq expression levels; genes with zero expression were excluded. **e,** Average H3K23me3 profiles corresponding to the gene groups shown in (**d**). **f,** RNA-seq analysis of *set-19(ok1813)* relative to wild-type worms. Volcano plot showing log_2_FoldChange (*set-19*/wt) versus -log_10_(adjusted p value [padj]), calculated using DESeq2. Significantly upregulated and downregulated genes are highlighted (padj < 0.01; |log_2_FoldChange| > 1). *N* = 3 biological replicates. **g,** H3K23me3 enrichment [log₂(IP/Input)] across gene bodies for genes upregulated in *set-19* mutants. Heatmaps showing a global reduction in the H3K23me3 signal in *set-19(ok1813)* relative to wild-type animals, with average enrichment profiles shown above.

To examine whether SET-19 is required for H3K23me3 occupancy across the genome, we compared H3K23me3 ChIP-seq profiles between wild-type and *set-19* mutant animals. The loss of *set-19* led to a substantial but partial reduction in H3K23me3 levels across the genome (Fig. 3b, c and Fig. S5c). Differential binding analysis using DiffBind ^56^ revealed that 55.4% of H3K23me3 peaks were significantly reduced in the *set-19* mutant relative to wild-type animals (Fig. S5d). Notably, the reduction in H3K23me3 was not uniform across chromosomal regions in *set-19* mutants, with substantial residual signals in certain chromosome arms and an apparent local increase near the tip of chromosome X (Fig. 3b and Fig. S5c). These patterns may reflect contributions from additional H3K23 methyltransferases or differences in local chromatin environments, although potential technical effects cannot be excluded.

H3K23me3 has been shown to be required for transcriptional repression of nuclear RNAi-targeted genes ^25,26^. To investigate the contribution of SET-19-dependent H3K23me3 to gene regulation, we performed mRNA-seq analysis in wild-type N2 and *set-19* mutant animals. Genes were grouped into five expression classes based on their expression levels in wild-type N2 animals, ranging from Class I (highest expression) to Class V (lowest expression), and then H3K23me3 levels were quantified for each class. H3K23me3 levels showed a strong negative correlation with gene expression, with lowly expressed genes exhibiting higher H3K23me3 occupancy (Fig. 3d, e). Transcriptome analysis revealed that 520 genes were significantly upregulated and 130 genes were downregulated in the *set-19* mutant compared with wild-type animals (Fig. 3f). The upregulated genes displayed reduced H3K23me3 levels in the *set-19* mutant (Fig. 3g). For example, *ule-1* and *col-8* showed reduced H3K23me3 levels accompanied by increased gene expression (Fig. S5e). *T05B9.1* and *asp-17* retained H3K23me3 enrichment and showed little or no change in RNA levels (Fig. S5f). These observations indicate that the loss of SET-19-dependent H3K23me3 is associated with transcriptional derepression at a subset of genes.

Interestingly, some highly expressed genes also exhibited relatively high H3K23me3 levels. We compared their chromatin features with those of genes with lower H3K23me3 levels within each expression class. H3K23me3-high genes showed higher H3K9me3 and H3K27me3, but lower H3K4me3 and H3K36me3, across expression classes (Fig. S6). Thus, even among highly expressed genes, relatively high H3K23me3 was generally associated with additional features of repressive chromatin rather than with a uniformly active chromatin state.

Because H3K23me3-high genes were also enriched for H3K9me3 and H3K27me3, we next asked whether the loss of SET-19-dependent H3K23me3 was accompanied by changes in these other marks. The genome-wide distributions of both marks were broadly similar between wild-type and *set-19* mutant animals (Fig. S7a). Differential binding analysis using DiffBind revealed a predominantly decreasing pattern for H3K9me3, with 583 peaks showing significantly reduced signals and 18 showing significantly increased signals in the *set-19* mutant (Fig. S7b). In contrast, H3K27me3 was largely unchanged (Fig. S7c). We next examined whether changes in H3K9me3 or H3K27me3 accompanied H3K23me3 loss, including at regions associated with transcriptional upregulation. The 2,378 regions with significantly reduced H3K23me3 were defined as H3K23me3-loss regions (Fig. S5d). Of these, 79 were associated with upregulated genes and were designated H3K23me3-loss-upregulated regions. Among the H3K23me3-loss regions, H3K27me3 was largely unchanged as well (Fig. S7d), and none of the 79 H3K23me3-loss-upregulated regions showed significant changes in H3K27me3 (Fig. S7e). In contrast, H3K9me3 was reduced at 189 of the 2,378 H3K23me3-loss regions (8.0%) (Fig. S8a). Among the 79 H3K23me3-loss-upregulated regions, five regions also showed reduced H3K9me3 (Fig. S8b). For example, *clec-206* and *Y53F4B.23* showed reduced H3K9me3 accompanied by increased RNA-seq signals, whereas *deg-1* showed no detectable change in H3K9me3 despite increased RNA-seq signals (Fig. S8c). These observations indicate that reduced H3K9me3 accompanies H3K23me3 loss at only a subset of genomic regions, whereas H3K27me3 remains largely unchanged.

Collectively, these results indicate that SET-19 contributes to genomic H3K23me3 occupancy and loss of SET-19 is associated with derepression of a subset of genes. Although H3K23me3 was associated with H3K9me3 and H3K27me3 across expression classes, changes in H3K27me3 were rare following SET-19-dependent H3K23me3 loss, whereas reduced H3K9me3 was observed at a subset of genomic regions.

### SET-19 is dispensable for feeding RNAi targeting both germline and somatic genes

H3K23me3 and the putative H3K23 methyltransferase SET-32 have previously been implicated in transgenerational epigenetic inheritance (TEI) ^25,26,32,34,57^. We first tested whether SET-19 is required for feeding RNAi targeting several germline and somatic genes, including *pos-1, mex-3, dpy-11*, and *unc-15*. RNAi targeting *pos-1* or *mex-3* causes embryonic lethality and results in unhatched F1 embryos ^58-60^. RNAi targeting *dpy-11* or *unc-15* induces dumpy and paralysis phenotypes, respectively ^61-63^. However, neither *set-19* nor *set-32* mutants displayed detectable defects in these feeding RNAi responses (Fig. S9a–d). RDE-1 is an essential Argonaute protein required for feeding RNAi response ^64^. As a control, *rde-1* mutants were completely resistant to feeding RNAi. To increase the sensitivity of feeding RNAi assays, we next used an enhanced RNAi background ^62^. In *eri-1* mutants, RNAi targeting *dpy-11* resulted in a strongly enhanced dumpy phenotype in the progeny. RNAi targeting *lir-1* induced larval arrest, and targeting *dpy-13* induced a super dumpy phenotype ^62,65,66^; both phenotypes were abolished by *nrde-3* mutation. However, the *eri-1;set-19* mutant remained fully competent for feeding RNAi against *dpy-11*, *lir-1*, and *dpy-13* (Fig. S9e–g).

We then asked whether SET-19 also participates in the transgenerational epigenetic inheritance of RNAi. We employed a germline-expressed *pie-1p::gfp::h2b* reporter and performed feeding RNAi against *gfp* for one generation, after which animals were transferred to fresh plates without continued RNAi exposure. As expected, wild-type animals silenced *gfp* expression in the parental generation and maintained robust silencing in their F1 progeny. In *hrde-1*, *set-25*, and *set-32* mutants, the parental generation remained sensitive to feeding RNAi, but silencing inheritance was abolished in the F1 progeny (Fig. 4a, b). In contrast, *set-19* mutants retained effective *gfp* silencing in both the parental and F1 generations, comparable to that in control animals (Fig. 4a, b). Similarly, we tested whether SET-19 is required for the intergenerational inheritance of RNAi using a soma-expressed *sur-5p::gfp* reporter. Feeding dsRNA targeting *gfp* resulted in efficient silencing of GFP expression in both parental animals and F1 progeny in control animals (Fig. 4c, d). As expected, GFP expression was not silenced in the *rde-1* mutant, consistent with the essential role of RDE-1 in feeding RNAi responses. NRDE-3 is required for the intergenerational inheritance of RNAi targeting somatic genes ^62^. In *nrde-3* mutants, the somatic *sur-5p::gfp* was silenced in the parental generation, but this silencing was abolished in the F1 progeny. In *set-19* and *set-32* mutants, *sur-5p::gfp* remained silenced in both the parental generation and F1 progeny (Fig. 4c, d).

**Fig. 4.**
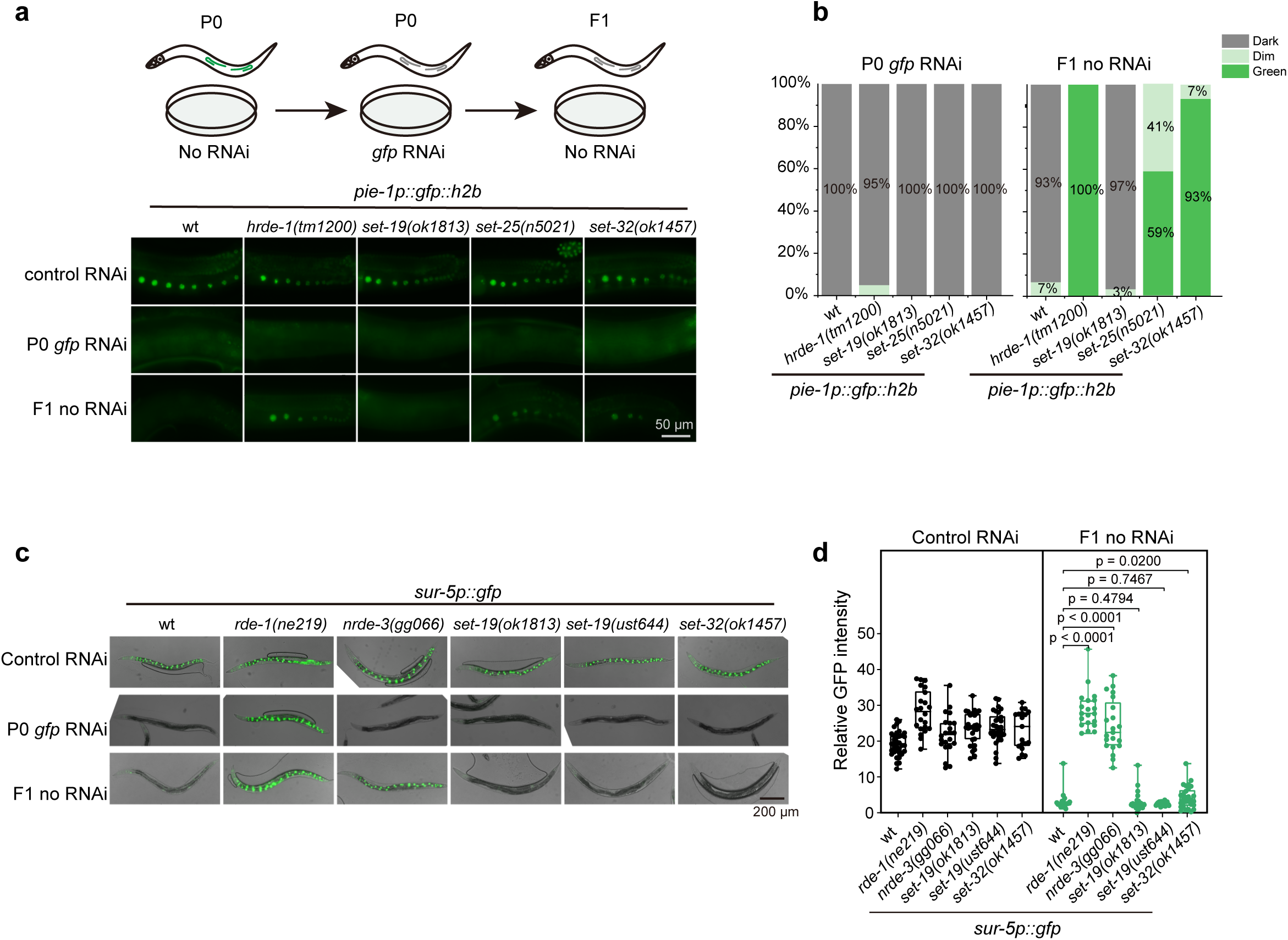
SET-19 is dispensable for feeding RNAi and its inheritance. **a,** Representative fluorescence images of animals expressing *pie-1p::gfp::h2b* subjected to *gfp* RNAi or control treatment. F1 progeny were maintained on control bacteria without feeding RNAi. **b,** The percentages of the indicated P0 and F1 animals expressing GFP were quantified. At least 30 animals were scored per genotype and generation. **c,** Representative fluorescence images of animals expressing *sur-5p::gfp* following *gfp* RNAi treatment. **d,** Quantification of relative GFP fluorescence intensity for animals shown in (**c**). Data are presented as the mean ± SD (*N* = 3 biological replicates; n > 18 worms per genotype). Statistical significance was assessed using a two-tailed t test.

Thus, these results suggest that SET-19 is dispensable for RNAi and its inheritance in *C. elegans*.

### SET-19 is predominantly expressed in somatic cells and required for developmental timing

To explore the physiological roles of SET-19, we examined the development and reproduction of *set-19* mutants, but did not observe significant defects in brood size or hatch rate compared with wild-type animals (Fig. 5a, b). However, *set-19* mutants exhibited a modest developmental delay relative to wild-type animals (Fig. 5c).

**Fig. 5.**
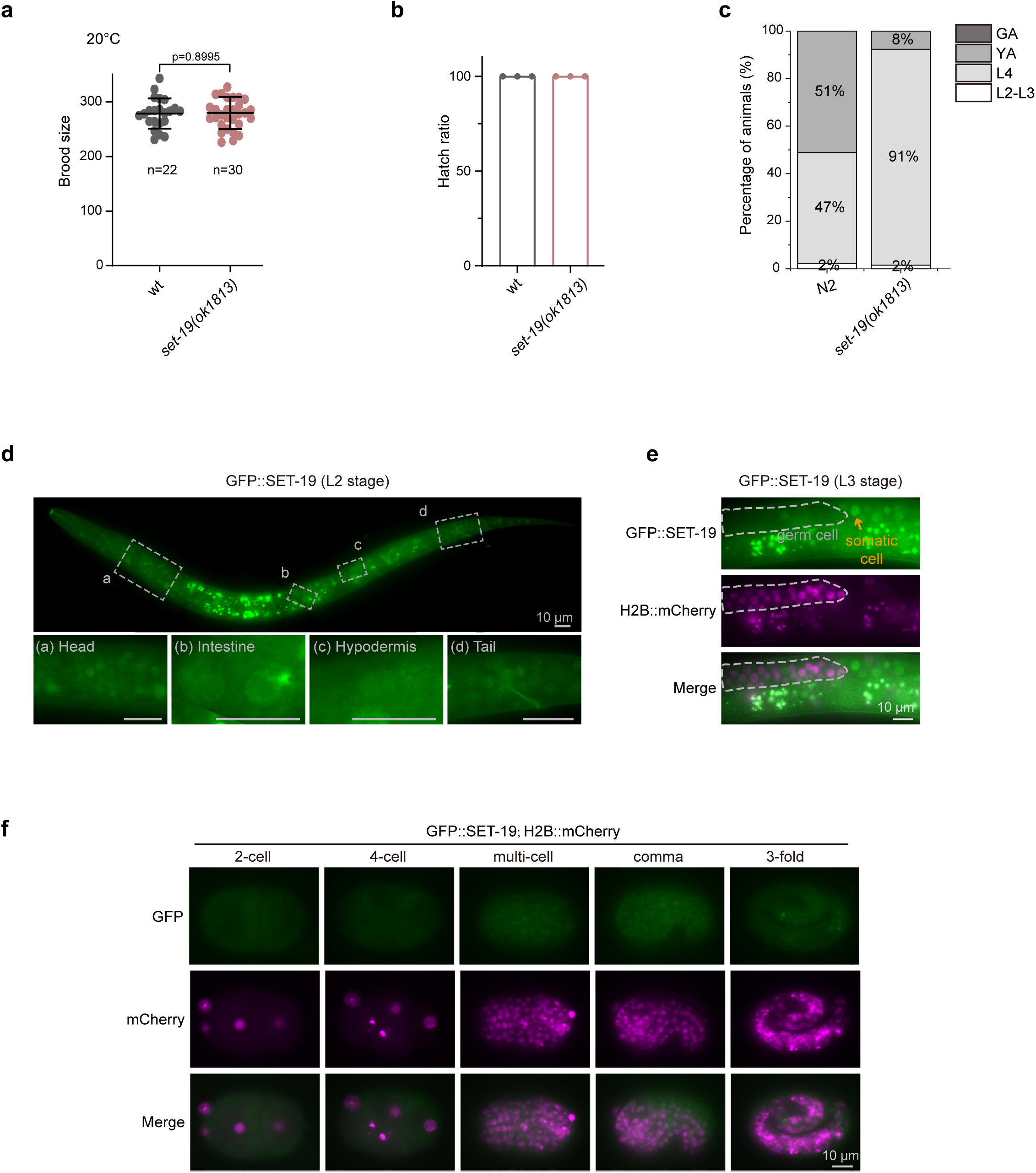
SET-19 is predominantly expressed in somatic cells and required for developmental timing. **a,** Brood size of the indicated genotypes maintained at 20°C. Individual L4 animals were transferred to fresh NGM plates, and total progeny were counted. **b,** Hatch rate of embryos laid by synchronized adult hermaphrodites at 20°C. The hatch rate was calculated as the ratio of hatched larvae to total embryos. N = 3 independent batches; wild type, n = 269; *set-19(ok1813)*, n = 227. **c,** Distribution of synchronized animals across developmental stages at 64 hours after egg laying at 20°C. The percentage of animals at each developmental stage was determined (n > 50 animals per genotype). L2–L3, second- to third-stage larvae; L4, fourth-stage larvae; YA, young adults; GA, gravid adults. **d,** Fluorescence images of L2 animals expressing GFP::SET-19. **e,** Fluorescence images of L3 animals expressing GFP::SET-19 and H2B::mCherry. GFP::SET-19 shows enrichment in somatic cells. **f,** Subcellular localization of GFP::SET-19 and H2B::mCherry in embryos.

Publicly available expression data from WormBase (version WS297) showed that *set-19* mRNA is more highly expressed in embryos than in larvae (Fig. S10a). To further examine cell type-specific expression, we analyzed published single-cell RNA-seq data from L2-stage larvae ^67^. *set-19* expression was relatively enriched in intestinal cells and neurons (Fig. S10b).

To directly assess the expression and subcellular localization of SET-19, we generated an endogenous GFP-tagged SET-19 strain using CRISPR/Cas9-mediated genome editing. GFP::SET-19 was broadly expressed in somatic cells during both embryonic and larval stages and was predominantly localized to the nucleus (Fig. 5d–f). During embryogenesis, SET-19 expression was low in early embryos but markedly increased in late embryos (Fig. 5f). No detectable GFP::SET-19 signal was observed in germ cells (Fig. 5e). In contrast, SET-32 was highly expressed in germ cells, displayed both nuclear and cytoplasmic localization during larval stages, and was weakly expressed in embryos (Fig. S10c–e).

These results indicate that SET-19 is predominantly expressed in somatic cells and is largely excluded from the germline.

### SET-19 conducts H3K23 trimethylation predominantly in somatic cells

To further examine the role of SET-19 in H3K23me3 in vivo, we performed immunofluorescence staining and assessed H3K23me3 levels across different developmental stages and tissues. In wild-type N2 animals, H3K23me3 signals were readily detected throughout development and were broadly distributed across tissues (Fig. 6a–d). Consistent with the somatic expression pattern of SET-19, the loss of *set-19* resulted in a pronounced reduction in H3K23me3 in somatic cells (Fig. 6a). However, no obvious reduction of H3K23me3 was observed in embryos (Fig. 6b). In young adult worms, the reduction in H3K23me3 was most prominent in intestinal cells (Fig. 6c), a tissue in which *set-19* expression is relatively enriched (Fig. S10b). Notably, H3K23me3 levels in germ cells remained largely unchanged in *set-19* mutants (Fig. 6d), consistent with the absence of detectable *set-19* expression in the germline (Fig. 5e). The limited changes in H3K23me3 levels detected by western blotting and immunofluorescence staining in *set-32* single mutant (Fig. 1e, Fig. 6a–d, and Figs. S2 and S3) do not exclude contributions from SET-32 in other cellular contexts or potential functional redundancy with other H3K23 methyltransferases.

**Fig. 6.**
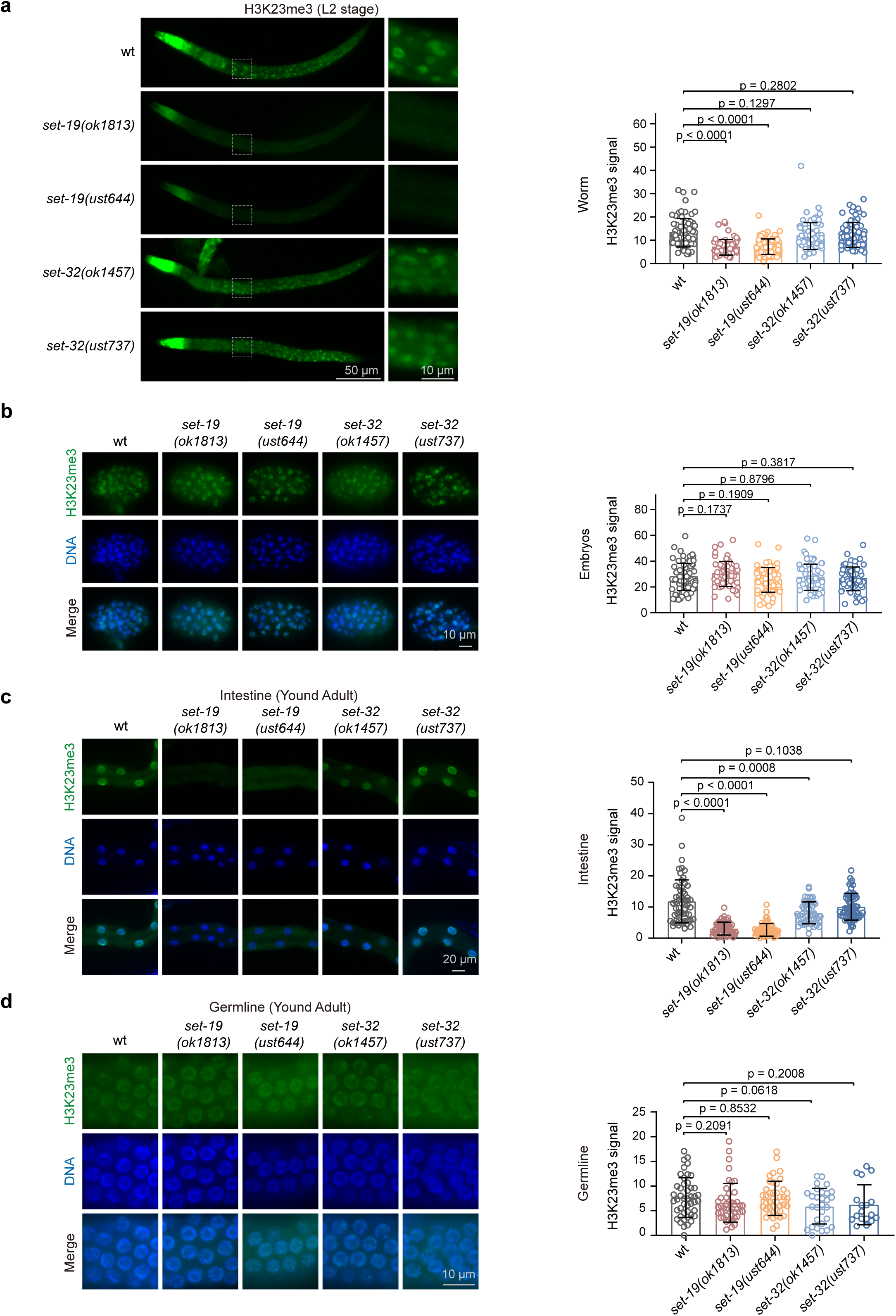
SET-19 mediates H3K23me3 predominantly in somatic cells. **a,** Representative immunofluorescence images of H3K23me3 staining in L2-stage worms (left) and quantification of whole-worm H3K23me3 fluorescence intensity (right). H3K23me3 levels were calculated as whole-worm fluorescence intensity after background subtraction. For L2-stage larvae: N = 3 independent batches; wild type, n = 78 worms; *set-19(ok1813)*, n = 64 worms; *set-19(ust644)*, n = 82 worms; *set-32(ok1457)*, n = 75 worms; *set-32(ust737)*, n = 77 worms. **b,** Representative H3K23me3 immunostaining in embryos (left) and quantification of H3K23me3 immunostaining in stage-matched embryos (right). H3K23me3 levels were calculated as nuclear fluorescence intensity after background subtraction, with three nuclei averaged per embryo. For embryos: N = 2 biological replicates; wild type, n = 78 embryos; *set-19(ok1813)*, n = 62 embryos; *set-19(ust644)*, n = 70 embryos; *set-32(ok1457)*, n = 65 embryos; *set-32(ust737)*, n = 65 embryos. **c,** Representative images (left) and quantification of H3K23me3 immunostaining in the intestinal nuclei of young adult worms (right). H3K23me3 levels were calculated as nuclear fluorescence intensity after background subtraction, with three nuclei averaged per worm. H3K23me3 (green); Hoechst 33342 (blue). For intestinal nuclei: wild type, n *=* 60 worms, *N =* 3 biological replicates; *set-19(ok1813)*, n *=* 60 worms, *N =* 3 biological replicates; *set-19(ust644)*, n *=* 60 worms, *N =* 3 biological replicates; *set-32(ok1457)*, n *=* 51 worms, *N =* 2 biological replicates; *set-32(ust737)*, n *=* 60 worms, *N =* 3 biological replicates. **d,** Representative images (left) and quantification of H3K23me3 immunostaining in the germline nuclei of young adult worms (right). H3K23me3 levels were calculated as nuclear fluorescence intensity after background subtraction, with three nuclei averaged per worm. H3K23me3 (green); DAPI (blue). For germline nuclei: wild type, *n =* 46 worms, *N =* 3 biological replicates; *set-19(ok1813)*, *n =* 44 worms, *N =* 3 biological replicates; *set-19(ust644)*, *n =* 42 worms, *N =* 2 biological replicates; *set-32(ok1457)*, *n =* 29 worms, *N =* 2 biological replicates; *set-32(ust737)*, *n =* 18 worms, *N =* 2 biological replicates. Signal quantification was performed using ImageJ. All data are presented as the mean ± SD. Two-tailed *t* tests were performed to determine statistical significance.

Together, these results support a major role for SET-19 in establishing H3K23me3 in somatic cells.

## Discussion

H3K23me3 is a histone modification presented across multiple species, yet its mechanisms of deposition and biological functions have remained poorly defined. In this study, we identified SET-19 as an H3K23me3 methyltransferase through a combination of biochemical and in vivo approaches. We show that SET-19-dependent H3K23me3 is enriched at heterochromatin-associated regions and is associated with transcriptional repression of a subset of genes. SET-19-dependent deposition of H3K23me3 occurs predominantly in somatic tissues in *C. elegans*. However, since our transcriptomic analysis was performed using whole worms, this work may not fully capture tissue-specific changes in gene expression. Future studies using tissue-specific transcriptomic approaches will further define the contribution of SET-19-dependent H3K23me3 to gene regulation in different cell types. Collectively, these findings extend our understanding of how H3K23 methylation is established and suggest that its deposition is regulated in a tissue-specific manner.

Histone lysine methylation is generally catalyzed by SET domain-containing methyltransferases. The *C. elegans* genome encodes at least 38 SET domain-containing proteins, yet only a subset of them have been functionally characterized ^37-46,68-71^. To date, efforts to identify the enzymes responsible for H3K23 methylation have been relatively limited. Candidate-based analyses assessed H3K23me2 by immunostaining in primordial germ cells and the adult germline, but failed to identify a bona fide KMT for germline H3K23me2 ^18^. SET-32 was then shown to methylate H3K23 in vitro and to be required for nuclear RNAi-induced H3K23me3 in vivo ^25,26^. Subsequently, the closely related methyltransferase SET-21 was also shown to possess H3K23 methyltransferase activity in vitro, predominantly generating H3K23me1 and H3K23me2. SET-32 and SET-21 are more strongly associated with the germline ^26^. Loss of *set-32* or *set-21* alone did not reveal a substantial reduction in H3K23 methylation, consistent with tissue-, stage-, or context-restricted functions and potential redundancy ^25,26,35^. Here, we show that SET-19 functions as a new H3K23me3 methyltransferase that acts primarily in somatic tissues. Based on our findings together with previous reports, we therefore propose a working model in which SET-19 predominantly mediates H3K23me3 deposition in somatic cells, whereas germline H3K23 methylation may rely more heavily on SET-32 and SET-21 (Fig. 7).

**Fig. 7.**
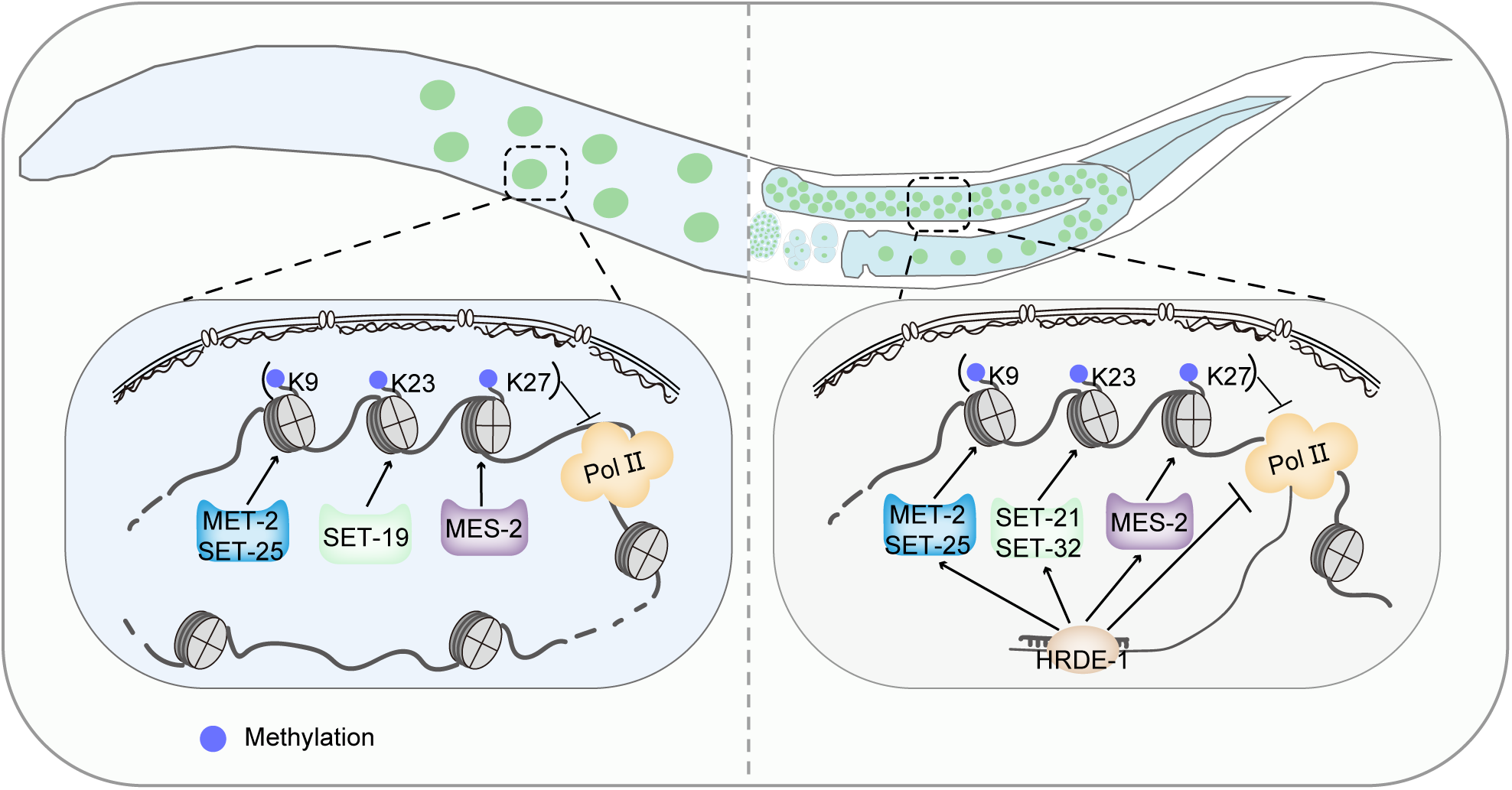
A working model for somatic and germline H3K23 methylation in *C. elegans*. (Left) In somatic cells, SET-19 acts as the major H3K23me3 methyltransferase. SET-19-catalyzed H3K23me3 coexists with H3K9me3 and H3K27me3 in heterochromatin-associated regions and is associated with transcriptional repression at a subset of genes. (Right) In the germ line, SET-32 and SET-21 are proposed to contribute to H3K23me3 deposition downstream of the HRDE-1-mediated RNAi pathway and to the transgenerational inheritance of RNAi.

Classical heterochromatin marks, such as H3K9 and H3K27 methylation, play essential roles in transcriptional repression, perinuclear anchoring of chromatin, genome stability, and RNAi-mediated gene silencing ^27-30,72-74^. Previous studies linked H3K23me3, particularly SET-32-associated H3K23 methylation, to nuclear RNAi and transgenerational epigenetic inheritance ^25,26,29,32,34,75^. Immunofluorescence staining shows that H3K23me3 forms discrete foci within the nucleus, resembling the nuclear distribution patterns reported for H3K9 and H3K27 methylation ^76^. In addition, the heterochromatin protein HPL-1, a chromodomain-containing reader, has been shown to recognize both H3K9 and H3K23 methylation ^18^. Our data show that H3K23me3 is enriched at heterochromatin-associated regions and is linked to transcriptional repression of a subset of genes. These observations suggest that H3K23me3 is an additional heterochromatin-associated repressive mark. However, H3K23me3-high genes in the highly expressed class were also enriched for H3K9me3 and H3K27me3, indicating that high transcript abundance can coexist with multiple repressive chromatin features. This pattern may reflect either tissue-specific differences obscured by whole-animal measurements or specialized transcriptional mechanisms operating within repressive chromatin environments, as exemplified by piRNA transcription from H3K27me3-enriched heterochromatic clusters in *C. elegans* ^77-79^. Together, these considerations suggest that H3K23me3 may contribute to transcriptional repression in specific genomic or cellular contexts. Nevertheless, whether H3K23 methylation contributes to perinuclear chromatin organization remains an intriguing question. Future studies using perinuclear tethering assays could be used to assess the contribution of H3K23 methylation to nuclear positioning ^76^.

H3K23 methylation appears to be functionally distinct from other heterochromatin marks. First, SET-19-dependent H3K23me3 loss was generally not accompanied by changes in H3K27me3 and was accompanied by reduced H3K9me3 only at a subset of genomic regions. These findings suggest that the transcriptional regulation associated with SET-19-dependent H3K23me3 cannot be solely explained by broad changes in these canonical repressive marks and that H3K23me3 may represent a partially distinct layer of heterochromatin-associated regulation, while retaining a limited local association with H3K9me3. Second, the methyltransferases responsible for these modifications differ in their tissue distributions and modes of regulation. The major H3K9 and H3K27 methyltransferases in *C. elegans*, MET-2/SET-25 and MES-2, respectively, act broadly across tissues and developmental stages ^44,80^. In contrast, SET-32 is expressed throughout the adult germline, whereas SET-21 is predominantly enriched in oocyte nuclei, and both proteins are also detected in embryos ^25,26^. Potential functional redundancy among these H3K23 methyltransferases could be tested directly by generating *set-32;set-21* and *set-19;set-32* double mutants. SET-19 contributes to H3K23me3 predominantly in somatic cells, pointing to distinct modes of enzymatic regulation among repressive histone modifications. Nevertheless, these distinct expression patterns do not exclude functional overlap among H3K23 methyltransferases, as SET-32 and SET-21 have been shown to function synergistically in H3K23me3 deposition at germline nuclear RNAi targets ^25,26,35^. Thus, different H3K23 methyltransferases may act redundantly under specific cellular conditions or contribute to multiprotein complexes responsible for H3K23 methylation. Although SET-32 has been linked to nuclear RNAi and transgenerational epigenetic inheritance in the germline ^25,26,29,32,34,75^, *set-19* mutants do not exhibit noticeable defects in feeding RNAi targeting either germline or somatic genes. These results indicate that SET-19 is not an indispensable requirement for the execution of feeding RNAi and its inheritance. However, whether SET-19 contributes to RNAi-induced H3K23me3 deposition in somatic cells requires further investigation.

While our findings support a predominantly somatic role for SET-19, the functions of H3K23 methylation in both somatic and germline cells remain to be fully defined. The unresolved questions include whether multiple H3K23 methyltransferases act redundantly or in parallel in the germline. It also remains unclear how SET-19-dependent and germline-enriched H3K23 methyltransferases are coordinated across tissues, and to what extent H3K23me3 overlaps functionally with other heterochromatin machineries. Systematic biochemical and genetic analyses of SET domain proteins in *C. elegans* will help define the full complement of histone methylation activities and clarify how chromatin-based regulation is directed to distinct developmental stages and cell types.

## Materials and Methods

### Strains

The Bristol strain N2 was used as the standard wild-type strain. All strains were grown at 20°C unless otherwise specified. The strains used in this study are listed in **Table S1**.

### Construction of plasmids and transgenic strains

For in situ expression of GFP::SET-19, the coding sequence of 3×FLAG::GFP, fused to a linker sequence (GGAGGTGGAGGTGGAGCT), was inserted immediately downstream of the start codon of *set-19* using the CRISPR/Cas9 system. The 3×FLAG::GFP fragment was PCR-amplified from YY178 genomic DNA. Homologous left and right arms (1.5 kb each) were amplified from N2 genomic DNA, and the backbone fragment was amplified from plasmid pCFJ151. All fragments were assembled into the repair plasmid using the ClonExpress MultiS One Step Cloning Kit (C113-02, Vazyme) via Gibson assembly. The injection mixture contained pDD162 (50 ng/µl), the repair plasmid (50 ng/µl), pCFJ90 (5 ng/µl), and two or three sgRNA plasmids targeting sequences near the N-terminus of the *set-19* gene (20 ng/µl per sgRNA plasmid). The mixture was injected into adult animals. Three to four days after injection, F1 worms expressing pharyngeal GFP were isolated using a Leica M165 FC fluorescence stereomicroscope. Progeny from these F1 worms were subsequently screened by PCR and validated by Sanger sequencing. The sequences of primers used for construction of the in situ transgenic strain are listed in **Table S2**.

### Construction of mutant strains via CRISPR/Cas9 technology

To generate sgRNA expression vectors, the 20 bp *unc-119* sgRNA guide sequence within the *pU6::unc-119* sgRNA(F+E) vector was replaced with distinct target-specific sgRNA guide sequences. A plasmid mixture containing 30 ng/µl of each of the three or four sgRNA expression vectors, 50 ng/µl of pDD162 plasmid, and 5 ng/µl of pSG259 was co-injected into wild-type N2 nematodes. Worms carrying the desired gene deletions were screened via PCR following the method described previously ^81^. All sgRNA sequences used in this study are listed in **Table S3**. The *set-19(ok1813)* and *set-32(ok1457)* deletion strains were obtained from the Caenorhabditis Genetics Center (CGC). The genotypes of *set-19(ok1813)*, *set-19(ust644)*, *set-32(ok1457)*, and *set-32(ust737)* were verified using two complementary PCR assays. Primer pairs flanking the deleted regions generated shorter PCR products from the corresponding mutant alleles than from wild type, whereas WT-specific primer pairs containing a primer within the deleted regions generated products from wild type but not from homozygous mutants. The *set-19(ust644)* and *set-32(ust737)* alleles generated in this study were further confirmed by Sanger sequencing of the deletion junctions. The primer sequences and detailed genotyping and validation information are provided in **Table S4**.

### Isolation of histones and mass spectrometry

Synchronized L3–L4 larvae and embryos (obtained by bleaching gravid adults) were washed three times with 1× M9 buffer. All samples were flash-frozen in liquid nitrogen and pulverized into a fine powder using a ball mill (60 Hz). The frozen powder was resuspended in Nuclear Extraction (NE) Buffer (100 mM HEPES, pH 7.5; 50 mM NaCl; 1% Triton X-100; 0.1% sodium deoxycholate; 1 mM EDTA; 10% glycerol; plus protease inhibitors). The resulting suspension was centrifuged at low speed to remove worm debris. To remove soluble proteins, the clarified supernatant was further centrifuged at high speed (16,000 × g for 10 min), and the resulting supernatant was discarded. The insoluble pellet was resuspended in NE buffer. Histone isolation was performed using a high-salt extraction strategy ^82^. First, the insoluble pellet was further lysed in salt-free buffer to disrupt nuclear envelopes; histones were subsequently liberated from chromatin complexes via extraction with 2.5 M NaCl buffer. Histones were separated by SDS-PAGE, and the Coomassie blue-stained band containing histone H3 was cut from the gel. The gel slices underwent two rounds of propionylation with propionic anhydride to derivatize free and monomethylated lysine residues. Afterward, the samples were digested with trypsin, followed by an additional propionylation step to modify newly exposed N-terminal amines ^41,83^.

The eluted peptides were analyzed on an Easy-nLC 1000 system (Thermo Fisher) coupled to a Q Exactive mass spectrometer (Thermo Fisher) by LC-MS/MS. The acquired RAW data were analyzed using EpiProfile 2.1_Celegans, an updated version of EpiProfile 2.0 ^84^. For each histone modified peptide, the relative abundance (%RA) was calculated by dividing the area under the curve (AUC) of each modified peptide by the sum of the areas corresponding to all the observed forms of that peptide.

### Western blotting

Synchronized L3–L4 larvae and embryos obtained by bleaching gravid adults were washed three times with 1× M9 buffer. Samples were stored at - 80°C until use. The samples were suspended in 2× SDS loading buffer and heated in a metal bath at 95°C for 5–10 minutes. The suspensions were then centrifuged at 16,000 × g, and the supernatants were collected. Proteins were resolved by SDS-PAGE on gradient gels (15% separation gel, 5% spacer gel) and transferred to a Hybond-ECL membrane. After washing with 1× Tris-buffered saline with Tween-20 (TBST) buffer and blocking with 5% milk-TBST, the membrane was incubated overnight at 4°C with antibodies. The membrane was washed three times for 10 minutes each with 1× TBST and then incubated with secondary antibodies at room temperature for 2 hours. The membrane was washed thrice for 10 minutes with 1× TBST and then visualized. The following antibodies were used for western blotting: anti-H3 (Abcam, ab1791), 1:8000; anti-H3K4me3 (Abcam, ab8580), 1:800; anti-H3K9me3 (Millipore, 07-523), 1:3000; anti-H3K27me3 (Millipore, 07-449), 1:8000; anti-H3K36me3 (Abcam, ab9050), 1:1000; anti-H3K23me1 (Active Motif, 39388), 1:6000; anti-H3K23me2 (Active Motif, 39653), 1:2000; anti-H3K23me2 (Beyotime, AG3990), 1:2000; anti-H3K23me3 (Active Motif, 61499), 1:3000; and anti-H3K23me3 (Beyotime, AG3889), 1:1000. Secondary antibodies: HRP-labeled goat anti-rabbit IgG (H +L) (Abcam, ab205718), 1:15000.

### Recombinant protein expression and purification

The DNA segments encoding the SET domain of SET-19 (SET-19 SET; aa 37–318), the SET and coiled-coil domains of SET-19 (SET-19 SET+CC; aa 37–454) and the full-length protein of SET-32 were amplified from a cDNA library of wild-type strain N2 using PCR. Subsequently, the SET-19 SET, SET-19 SET+CC and SET-32 constructs were cloned and inserted into the pGEX-4T-1 plasmid and expressed in *E. coli* BL21-GOLD (DE3) cells (Novagen). The recombinant proteins were affinity-purified via GST tag binding to glutathione beads (Smart-Lifesciences, SA008100) according to the manufacturer’s instructions. The eluted protein was dialyzed against a buffer containing 20 mM Tris-HCl pH 7.5, 5% glycerol, and 200 mM NaCl and subsequently concentrated. The concentrated protein was then subjected to high-speed centrifugation; after centrifugation, the clarified protein solution was supplemented with glycerol and stored at -80°C.

### In vitro histone methylation assay

GST-fused proteins (0.25 µM) were incubated with 1 µM histone H3 (Active Motif, 31294) and 80 µM S-adenosylmethionine (SAM) (NEB, B9003S) in a 20 µl reaction mixture containing methyltransferase activity buffer (20 mM Tris-HCl, pH 8.0; 50 mM NaCl; 20 mM KCl; 10 mM MgCl₂; 5% glycerol; 0.02% Triton X-100; 0.1 mg/ml BSA; and 1 mM dithiothreitol [DTT]) for 3 h at 20°C. Reactions were terminated by the addition of SDS loading buffer, followed by boiling for 15 min. Proteins were resolved on a 15% SDS-PAGE gel and analyzed by western blotting.

### Imaging

For imaging larval stages, animals were immobilized in 0.1 M sodium azide and mounted on 1.5% agarose pads. For imaging embryos and germ cells, gravid adults were dissected on a coverslip in 2 μl of 0.4× M9 buffer containing 0.1 M sodium azide and then mounted on freshly prepared 1.1% agarose pads. Imaging was performed using a Leica THUNDER Imaging System equipped with a K5 sCMOS camera and HC PL FLUOTAR objectives (100×/1.40–0.70 oil, 40×/0.80, and 20×/0.80). Images were acquired using Leica Application Suite X software (v3.7.4.23463).

### Immunofluorescence staining and quantification

Embryos were freeze-cracked using liquid nitrogen, then fixed in methanol for 30 seconds and post-fixed in 1% paraformaldehyde (PFA) for 2 minutes. After three washes in PBS containing 0.25% Triton X-100 (PBS-T), samples were permeabilized in PBS with 1% Triton X-100 for 20 minutes. Subsequently, the samples were blocked in PBS containing 0.5% BSA before overnight incubation with the primary antibody against H3K23me3 (Active Motif, 61499) at 4°C. Following three washes with PBS-T, samples were incubated with secondary antibodies (Alexa Fluor 488-conjugated anti-rabbit) for 2 hours at room temperature, before three final PBS-T washes and DNA staining with Hoechst 33342. For germline and intestine staining, gravid adult worms were dissected in M9 buffer, fixed in methanol for 1 minute at -20°C and in 2% PFA for 5 minutes. For L2 worm staining, worms were freeze-cracked using liquid nitrogen, then fixed in 1% paraformaldehyde (PFA) for 20 minutes. Fixed animals were permeabilized by sequential thiol-based reduction and oxidation steps. Samples were incubated in PBS containing 10% β-mercaptoethanol for 15 minutes, treated with PBS containing 10 mM DTT for 15 minutes, and then incubated in PBS containing 0.3% H₂O₂ for 15 minutes. The animals were subsequently fixed in methanol for 30 seconds and treated with the permeabilization solution (PBS containing 1% Triton X-100) for 20 minutes. Images were taken on a Leica THUNDER Imaging System with identical gain settings within each experimental set. Quantification was performed using ImageJ by measuring the mean gray value within the nuclear region and that of an adjacent background area. For each worm or tissue, fluorescence intensity was calculated as the average nuclear signal from three nuclei, subtracting the average background signal from three corresponding adjacent background regions. At least two independent experiments were performed.

### Brood Size

L4-stage hermaphrodites were singled onto plates and transferred daily as adults until embryo production ceased, and the progeny numbers were scored.

### Hatch ratio

Synchronized adult hermaphrodites were transferred to NGM plates to lay eggs, which were subsequently removed. The numbers of embryos and hatched larvae were scored, and the hatch ratio was calculated as the number of hatched larvae divided by the total number of embryos.

### Developmental stage assessment

Synchronized adult hermaphrodites were transferred to NGM plates to lay eggs, which were subsequently removed. The growth of the progeny was monitored every 24 hours at 20°C.

### RNAi

RNAi experiments were performed at 20°C by placing synchronized embryos on RNAi feeding plates as previously described ^85^. HT115 bacteria expressing the empty vector L4440 (a gift from A. Fire) were used as controls. Bacterial clones expressing target-specific double-stranded RNAs (dsRNAs) were obtained from the Ahringer RNAi library ^86^ and their identities were verified by Sanger sequencing.

### Total RNA isolation

Synchronized L4-stage worms were washed three times with 1× M9 to remove bacterial residues. For RNA extraction, 400 µl of TRIzol Reagent (Ambion, 15596026) was added to a 100 µl worm aliquot, followed by 7–8 cycles of freezing in liquid nitrogen and thawing in a 42°C water bath. Afterward, 100 µl of DNA/RNA Extraction Reagent (Solarbio LIFE SCIENCES, P1014) was added to the samples, which were then centrifuged at 16000 × g for 15 minutes at 4°C. The collected supernatant was further treated with 400 µl isopropanol and 400 µl pre-chilled 75% ethanol, and subjected to DNase I digestion (Thermo Fisher Scientific, EN0521). Finally, RNA was eluted into 20 µl of nuclease-free water and used for mRNA library preparation.

### mRNA-seq analysis

The Illumina-generated raw reads were first filtered to remove adapters, low-quality reads, and contaminants, yielding high-quality clean reads. The clean reads were aligned to the reference genome of WBcel235 via HISAT2 (v2.1.0) ^87^. For quantification, featureCounts (v1.6.0) ^88^ was used to calculate gene-level read counts. Library size factors were calculated using DESeq2 (v1.46.0) ^89^ with its standard median-of-ratios method based on genome-wide gene expression counts. Genes with an adjusted *P* value (padj) < 0.01 and an absolute log_2_FoldChange > 1 were considered significantly differentially expressed. Three biological replicates were analyzed per condition. All statistical analyses and plots were generated using custom R scripts.

### Chromatin immunoprecipitation (ChIP)

ChIP experiments were performed as previously described ^30^. For L3–L4 worms, frozen animals were first ground to a fine powder in liquid nitrogen ^90^. Then, the samples were crosslinked in 2% formaldehyde for 10 minutes at room temperature with gentle rotation. Crosslinking was quenched by adding 0.125 M glycine (final concentration) for 5 minutes at room temperature. Samples were sonicated for 13 cycles (30 s on and 30 s off per cycle) at high output with a Bioruptor Plus (Diagenode), with the sample tubes kept in an ice-water bath throughout the process. The lysates were precleared and immunoprecipitated with a rabbit anti-H3K23me3 antibody (Active Motif, 61499) overnight at 4°C. Chromatin/antibody complexes were recovered with Dynabeads^TM^ Protein A (Invitrogen, 10002D) followed by extensive sequential washes with 150 mM, 500 mM, and 1 M NaCl, respectively. Crosslinks were reversed overnight at 65°C. The recovered DNA was treated with RNase (Roche) for 30 minutes at 65°C, and all DNA samples were purified using a DNA purification kit (TianGen, #DP204).

H3K9me3 and H3K27me3 ChIP experiments were performed using the same procedure as described above for H3K23me3, with the antibodies listed in **Table S5**.

### ChIP-Seq

The DNA samples from the ChIP experiments were subjected to quality control (QC) prior to sequencing and then sequenced at high depth at Novogene Bioinformatics Technology Co., Ltd. (Beijing, China). Briefly, the ChIP DNA was processed for library construction following the standard protocol: the DNA fragments were combined with an End Repair/A-tailing mix and incubated in a single reaction to complete end repair and 3’-end adenylation. The resulting A-tailed DNA fragments were then incubated with sequencing adapters in ligation mix to achieve adapter ligation. The adapter-ligated DNA was subjected to size selection and several rounds of PCR amplification to obtain the final DNA library. The size distribution of the library fragments was analyzed on the Agilent 5400 system (Agilent, USA; with matching Agilent assay reagents). The library was quantified by qPCR to a final concentration of 1.5 nM. The qualified libraries were pooled according to their effective concentration and required data volume and then further amplified on cBot to generate clusters on the flow cell and sequenced with a paired-end 150 bp (PE150) method on an Illumina platform.

Published ChIP-seq datasets of histone modifications in *C. elegans* were downloaded from the NCBI GEO or ENCODE databases. The datasets used in this study are listed in **Table S5**.

### ChIP-seq data analysis

ChIP-seq data analysis was performed as previously described ^91^. ChIP-seq reads were aligned to the WBcel235 assembly of the *C. elegans* genome using Bowtie2 (v2.3.5.1) ^92^ by Ben Langmead with the default settings. The SAMtools (v0.1.19) ^93^ “view” utility was used to convert the alignments to BAM format, and the “sort” utility was used to sort the alignment files. ChIP-seq peaks were called using MACS2 (v2.1.1) ^94^ with subcommand. For the H3K4me3 and H3K36me3 datasets, which display narrow enrichment patterns, peaks were identified using the parameters “-g ce -B -f BAM -q 0.01”. For the H3K27me3, H3K9me3, and H3K23me3 datasets, which show broad histone modification domains, peaks were identified using the parameters “-g ce -B -f BAMPE --broad --broad-cutoff 0.01”. deepTools subcommand bamCoverage (v3.5.0) was used to produce bigWig tracks for data visualization with defined parameters (--binSize 20 --normalizeUsing BPM -- smoothLength 60 --extendReads 150) from bam files. The Integrative Genomics Viewer genome browser ^95^ was applied to visualize signals and peaks. The genomic regions were defined as chromosome arms and centers based on previous works ^53^. For visualization of peak locations, we used the R package ggplot2 (v4.0.0).

For heatmap analysis, the called peaks of H3K23me3 rep1 were used. deepTools subcommand computeMatrix (v3.4.3) was used to calculate the score matrix for the heatmap with defined parameters (reference-point --referencePoint center -b 3000 -a 3000 --skipZeros). The heatmap was plotted with deepTools subcommand plotHeatmap (v3.5.0).

### ChIPseqSpikeInFree normalization for H3K23me3

H3K23me3 ChIP-seq experiments were performed with two replicates each in both wild-type (wt) and *set-19* mutant backgrounds. To compare the H3K23me3 signals between the two backgrounds in the absence of spike-in, ChIPseqSpikeInFree (v1.2.4) ^96^ was used to calculate the scaling factor (SF) for every sample (SF_i_). The effective ChIP-seq library size for sample i was calculated as N_i_ ∗ SF_i_, where N_i_ is the original library size. The effective library size was then used to normalize the read count from sample i during the downstream differential analysis and heatmap visualization. For the two replicates of wt and *set-19* mutants, pairwise comparisons were performed, and the resulting SF values were averaged, yielding SF_wt = 1 and SF_set-19 = 2.115. The “-scale” parameter for bedtools genomecov was calculated from the effective library size using the formula: Scale parameter = 15000000/(Ni × SFi), where 15000000 is the preset reference read count. Normalized BEDGraph files were generated by setting the “-scale” parameter to the value from the above formula. The genome was then partitioned into fixed 20 bp windows, and the normalized signal was mapped into these windows using a bedtools map with weighting by the actual overlap length. The log₂(IP/Input) enrichment value for each window was subsequently calculated after adding a pseudocount of 1 to both IP and Input signals to avoid log₂(0) errors. Finally, the sorted BEDGraph file was converted into BigWig format for visualization. For the differential analysis of H3K23me3 peaks, we used the R package DiffBind (v3.16.0) ^56^ and integrated the sample-specific SF into the total read count normalization via the underlying DESeq2 (v1.46.0) ^89^ framework.

### Statistics

Bar graphs with error bars are presented with mean and standard deviation (SD). All experiments were conducted with independent *C. elegans* animals for the indicated N times. Statistical analysis was performed with a two-tailed Student’s t test.

## Acknowledgments

We are grateful to the members of the Guang laboratory for their comments and suggestions. We are grateful to the International *C. elegans* Gene Knockout Consortium and the National Bioresource Project for providing the strains. Some strains were provided by the CGC, which is funded by the NIH Office of Research Infrastructure Programs (P40 OD010440).

## Author contributions

Y.S., M. Huang, X.F. and S.G. conceptualized the research; M.X., X.F. and S.G. designed the research; M.X., Z.F., M. Huang, and C.Y. performed the research; X.C., X. Huang, C.Z., M. Hong, Z.Z., and X. Hou contributed new reagents; J.C., X. Hou, S.L. and M.L. provided technical guidance on bioinformatics analysis and protein purification; M.X., X.F. and S.G. wrote the paper.

## Data availability

The raw sequence data reported in this paper have been deposited in the Genome Sequence Archive in the National Genomics Data Center, China National Center for Bioinformation/Beijing Institute of Genomics, Chinese Academy of Sciences (GSA: CRA037051, CRA052996). The mass spectrometry proteomics data reported in this paper have been deposited in the iProX repository under accession codes IPX0015174000 and PXD073043.

## Figure legends

**Fig. S1.**
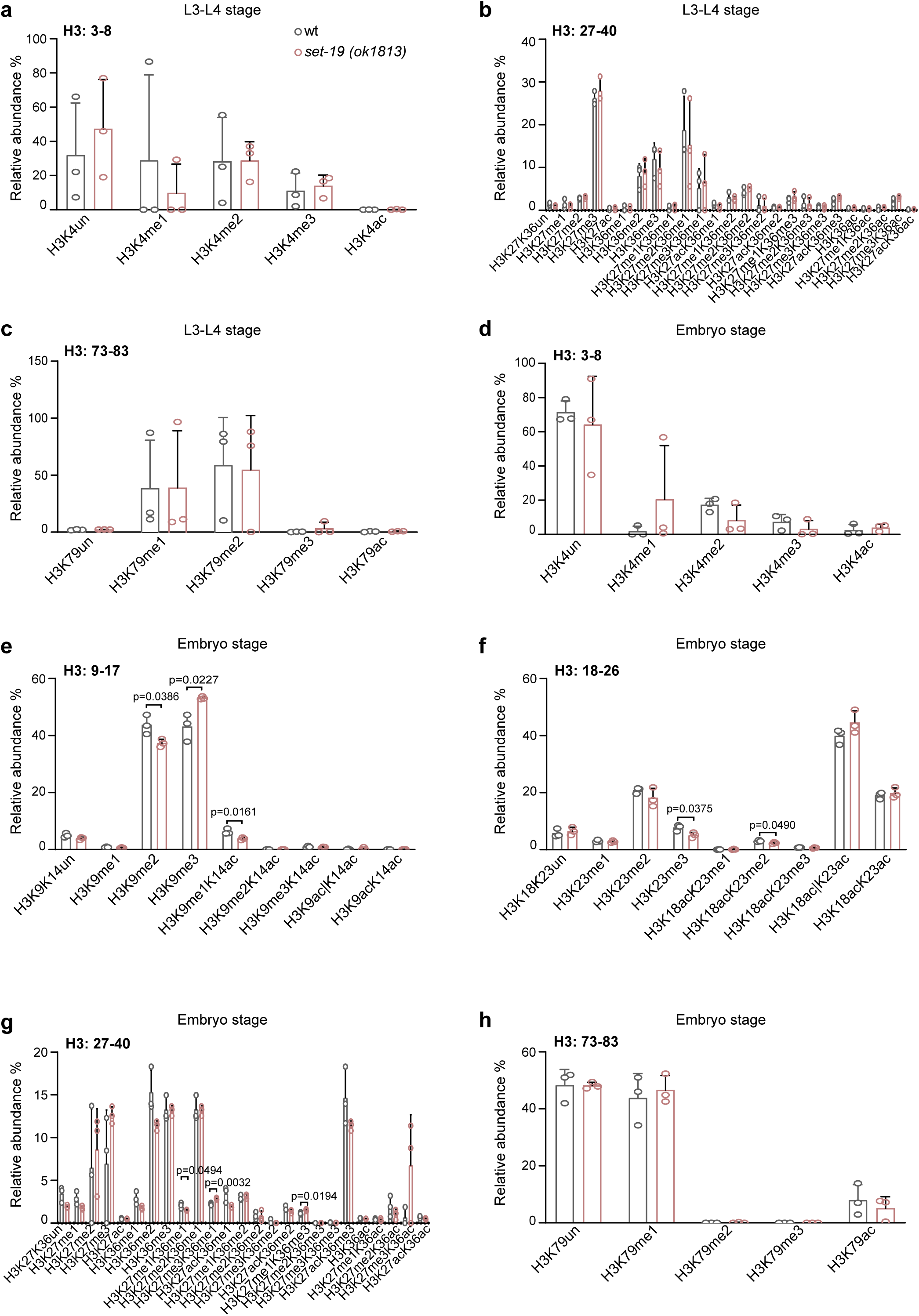
Mass spectrometry analysis of global H3 methylation in *set-19(ok1813)* mutants. Quantification of H3 methylation levels in wild-type and *set-19(ok1813)* mutants by quantitative mass spectrometry. H3 methylation levels were quantified as relative abundances, calculated by dividing the area under the curve (AUC) of each modified peptide by the sum of the AUCs of all observed forms of that peptide and multiplying by 100. Differences without indicated p values are not statistically significant (two-tailed t test, *p* < 0.05). *N =* 3 biological replicates.

**Fig. S2.**
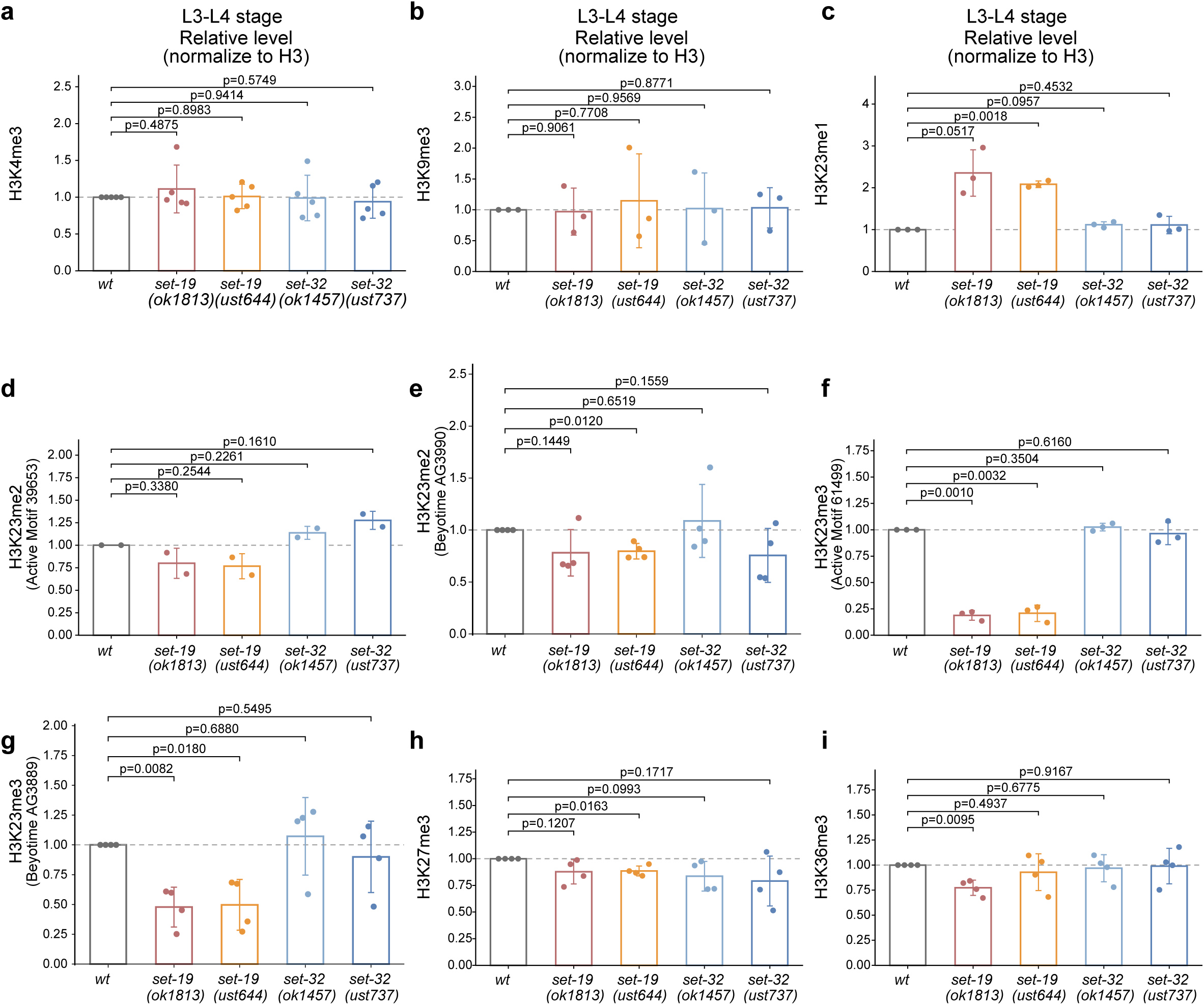
Quantification of histone H3 methylation levels in L3–L4-stage worms. **a–i,** Quantification of histone H3 methylation levels by western blotting in L3–L4-stage worms of the indicated genotypes, corresponding to **Fig. 1e**. H3K4me3 (**a**), H3K9me3 (**b**), H3K23me1 (**c**), H3K23me2 (**d, e**), H3K23me3 (**f, g**), H3K27me3 (**h**), and H3K36me3 (**i**) levels were quantified. Two independent antibodies were used to detect H3K23me2 (**d,** Active Motif; **e,** Beyotime) and H3K23me3 (**f,** Active Motif; **g,** Beyotime). Band intensities were normalized to H3 and expressed relative to wild-type (wt) controls. Data are presented as the mean ± SD, with individual points representing biological replicates. Statistical significance was assessed using a two-tailed t test. P values are indicated.

**Fig. S3.**
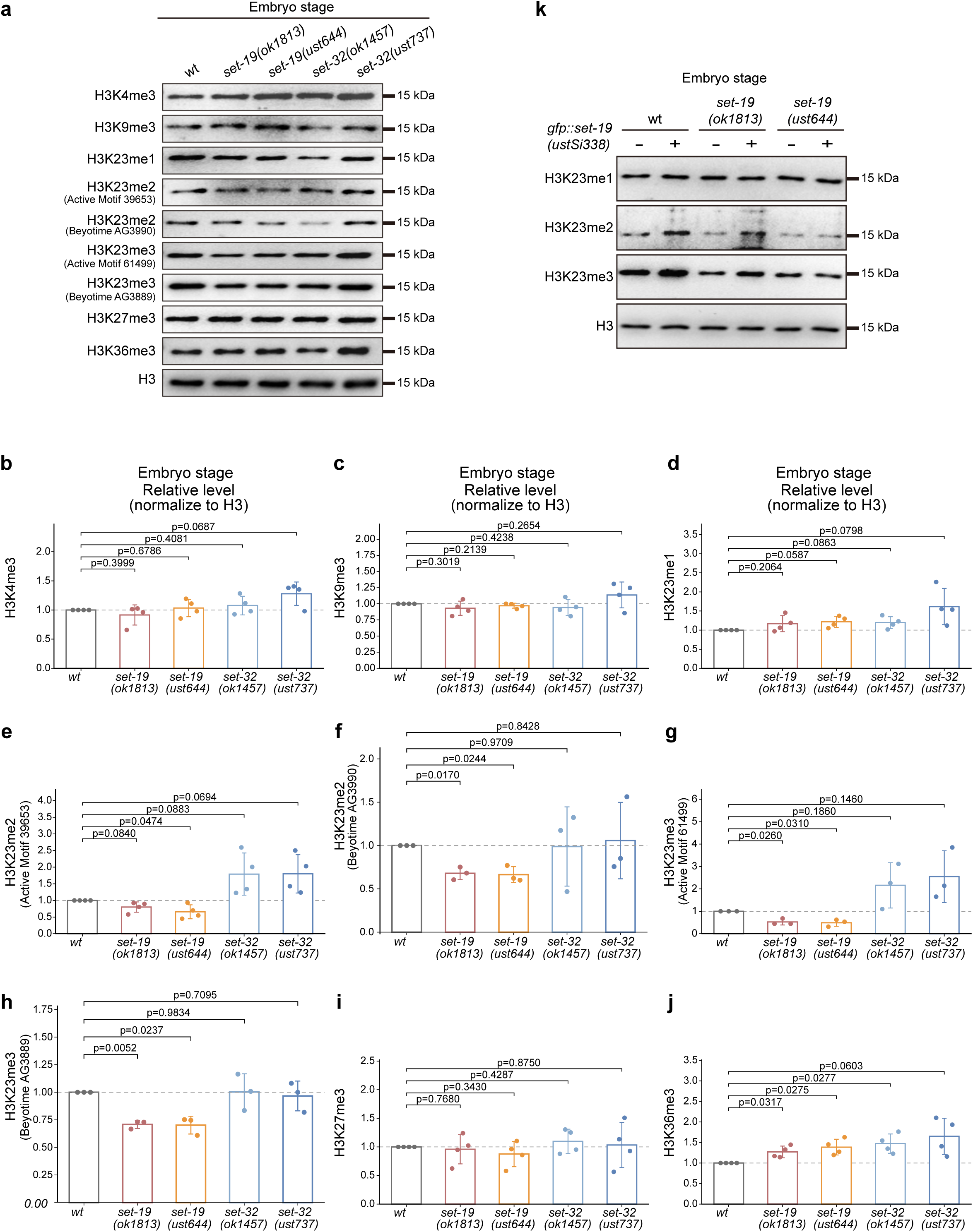
Western blotting analysis of histone H3 methylation levels in embryos. **a,** Representative western blotting analysis of histone H3 methylation levels in embryos of the indicated genotypes. **b – j,** Quantification of H3K4me3 (**b**), H3K9me3 (**c**), H3K23me1 (**d**), H3K23me2 (**e, f**), H3K23me3 (**g, h**), H3K27me3 (**i**), and H3K36me3 (**j**) levels in embryos. Two independent antibodies were used to detect H3K23me2 (**e,** Active Motif; **f,** Beyotime) and H3K23me3 (**g,** Active Motif; **h,** Beyotime). Band intensities were normalized to H3 and expressed relative to wild-type (wt) controls. Data are presented as mean ± SD, with individual points representing biological replicates. Statistical significance was assessed using a two-tailed t test. P values are indicated. **k,** Western blotting analysis of H3K23me1/2/3 levels in embryos. The *fib-1p::gfp::set-19(ustSi338)* transgene was used for rescue.

**Fig. S4.**
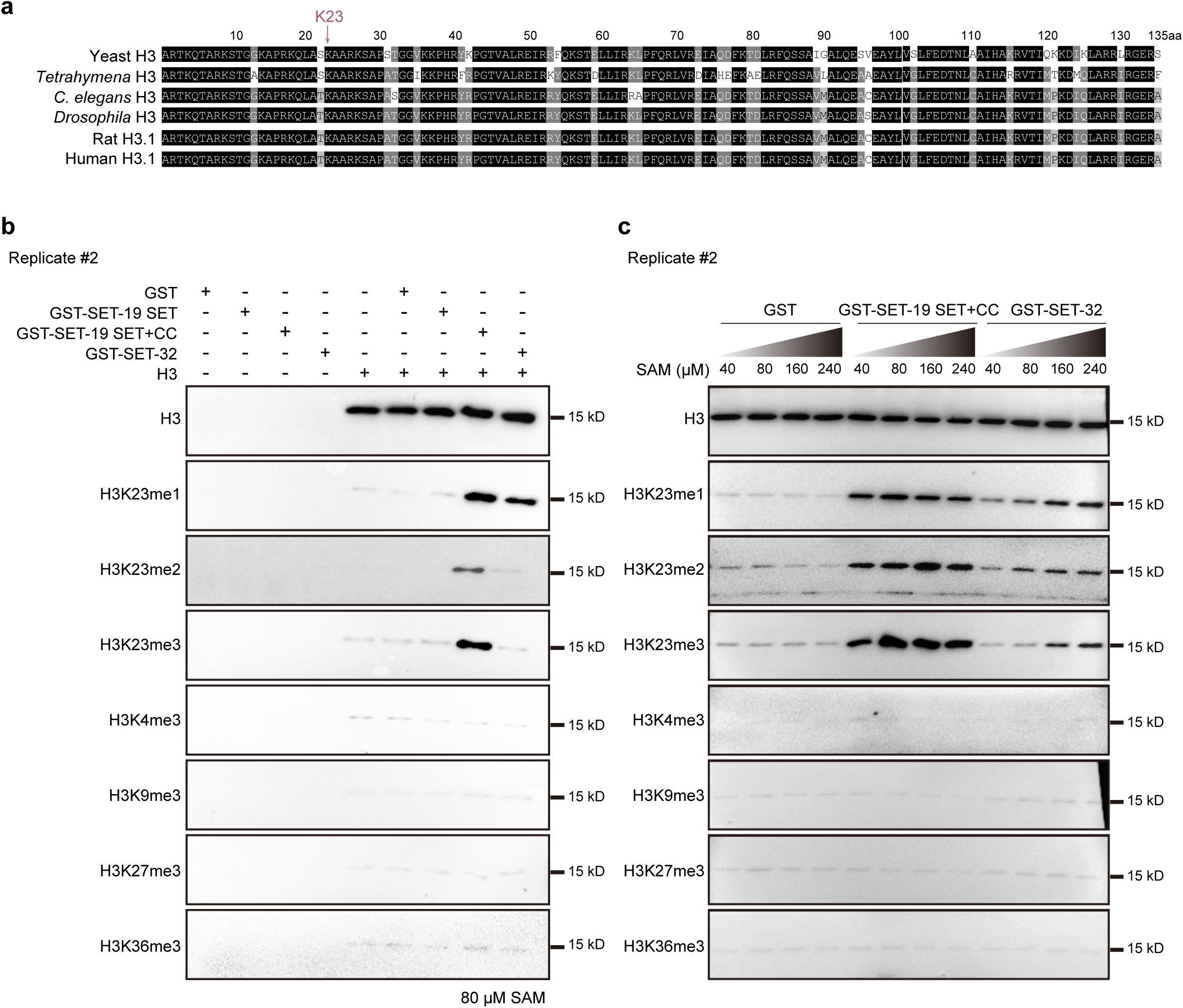
SET-19 specifically catalyzes histone H3K23 methylation in vitro. **a,** Multiple sequence alignment of histone H3 proteins across species, highlighting the conserved lysine 23 residue. **b, c,** Independent replicates of in vitro methylation assays corresponding to **Fig. 2b** and **Fig. 2c**, respectively.

**Fig. S5.**
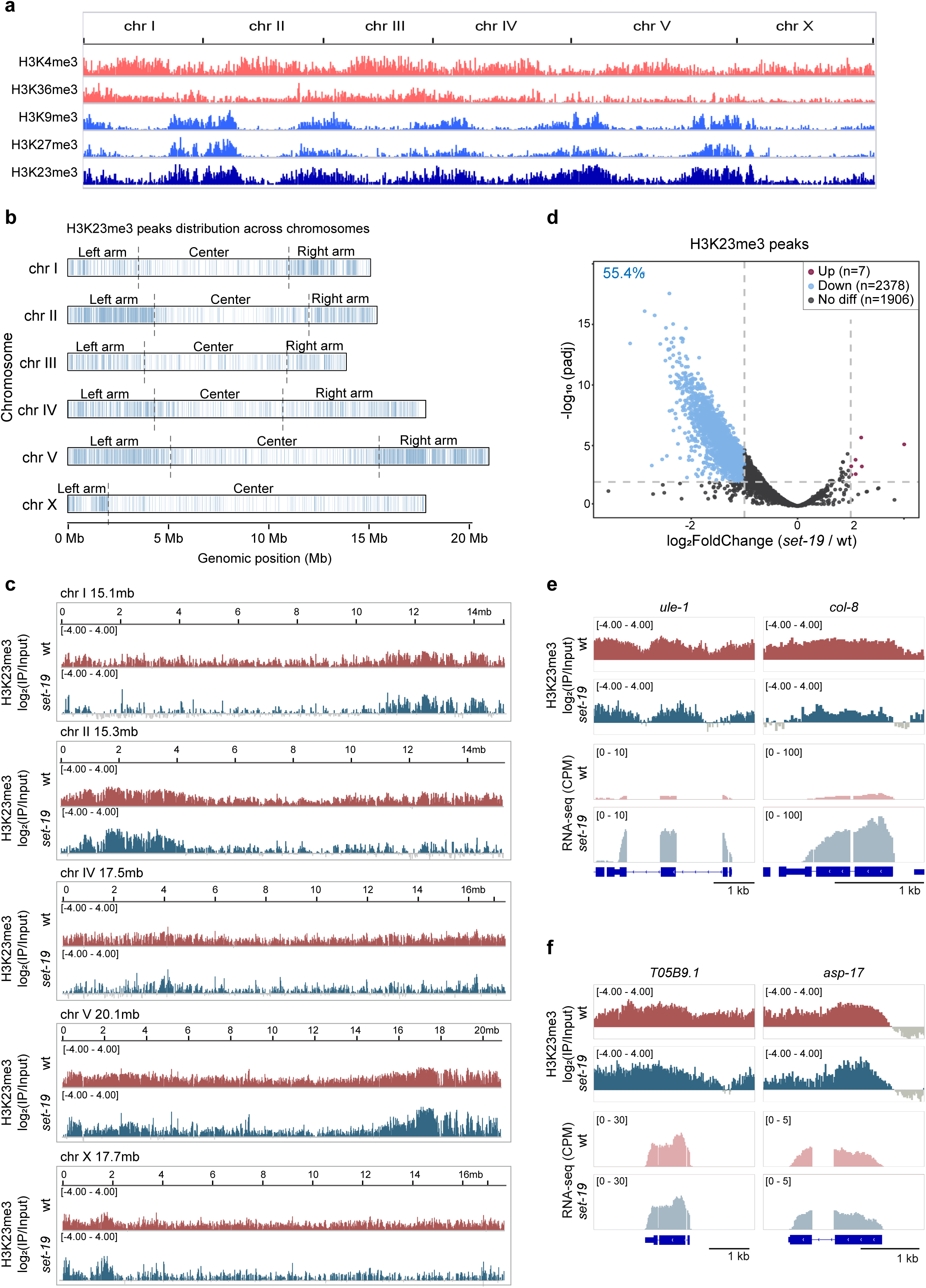
The loss of *set-19* alters genomic H3K23me3 occupancy. **a,** Genome-wide distribution of ChIP-seq peaks for H3K4me3, H3K36me3, H3K9me3, H3K27me3, and H3K23me3, as identified by MACS2. **b,** Distribution of H3K23me3 ChIP-seq peaks across chromosomes in wild-type worms. Peak positions identified by MACS2 from one representative biological replicate (replicate 1) are shown along each chromosome. Chromosome arms and centers are indicated. **c,** Comparison of H3K23me3 ChIP-seq signal distributions across chromosomes in wild-type and *set-19(ok1813)* worms. Mean log₂ enrichment over input is shown (*N* = 2 biological replicates). **d,** Differential H3K23me3 peak analysis between *set-19(ok1813)* and wild-type worms. Volcano plot showing log_2_FoldChange (*set-19*/wt) versus -log_10_(adjusted p value [padj]) from DiffBind ^56^ analysis. Differentially enriched peaks are highlighted (|log_2_FoldChange|>1 and padj < 0.01). *N* = 2 biological replicates. **e,** Representative genome browser tracks at *ule-1* and *col-8* showing reduced H3K23me3 enrichment accompanied by increased RNA-seq signals in *set-19* mutants compared with wild-type (wt) animals. H3K23me3 tracks show mean log_2_(IP/Input) enrichment from two biological replicates, whereas RNA-seq tracks show mean CPM-normalized coverage from three biological replicates. Signal ranges are indicated in brackets. Gene models are shown below the tracks. Scale bars, 1 kb. **f,** Representative genome browser tracks at *T05B9.1* and *asp-17* showing retained H3K23me3 enrichment and little or no change in RNA-seq signals in *set-19* mutants compared with wild-type animals. Track normalization and annotations are as described in (e).

**Fig. S6.**
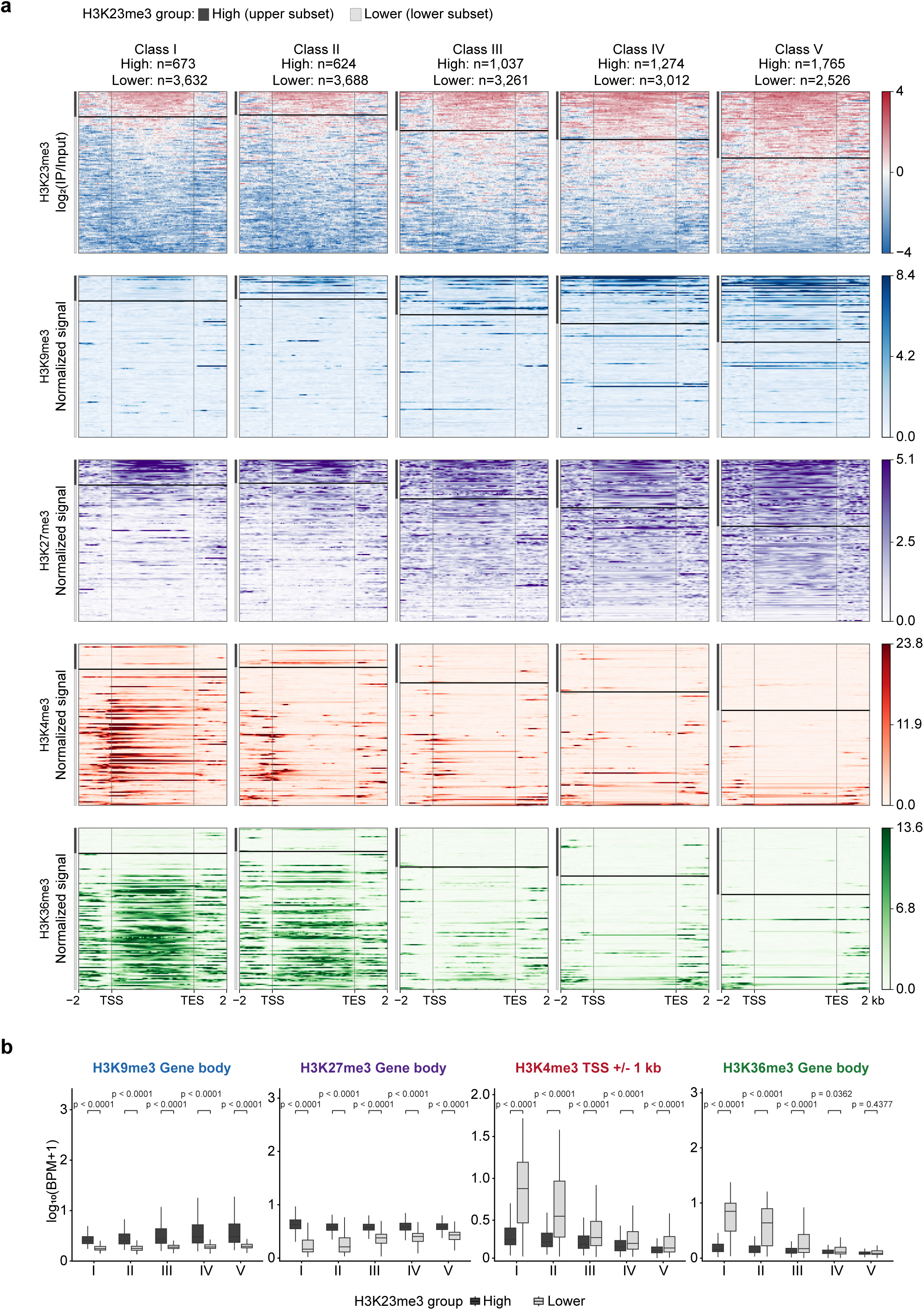
H3K23me3-high genes exhibit distinct chromatin features across expression classes. **a,** Heatmaps showing H3K23me3, H3K9me3, H3K27me3, H3K4me3, and H3K36me3 signals across genes grouped into five classes based on their expression levels in wild-type animals. For each gene, the mean H3K23me3 log₂(IP/Input) signal across the gene body was calculated. Among the 21,492 genes included in the analysis, the upper quartile was defined as H3K23me3-high and the remaining genes as H3K23me3-lower. Genes within each expression class were separated into H3K23me3-high and -lower groups and ordered by decreasing H3K23me3 signal. Heatmaps show the regions from 2 kb upstream of the transcription start site (TSS) to 2 kb downstream of the transcription end site (TES), with gene bodies scaled to 5 kb. The numbers of genes in each group are indicated. **b,** Quantification of normalized H3K9me3, H3K27me3, H3K4me3, and H3K36me3 ChIP-seq signals in the H3K23me3-high and -lower groups within each expression class. H3K9me3, H3K27me3, and H3K36me3 signals were averaged across gene bodies, whereas H3K4me3 signals were averaged within ±1 kb of the TSS. Values are shown as log₁₀(BPM + 1). Boxes indicate the median and interquartile range, and whiskers extend to 1.5 times the interquartile range; outliers are not shown. Statistical significance was assessed using a two-sided Wilcoxon rank-sum test, with Benjamini-Hochberg correction performed separately for each histone modification.

**Fig. S7.**
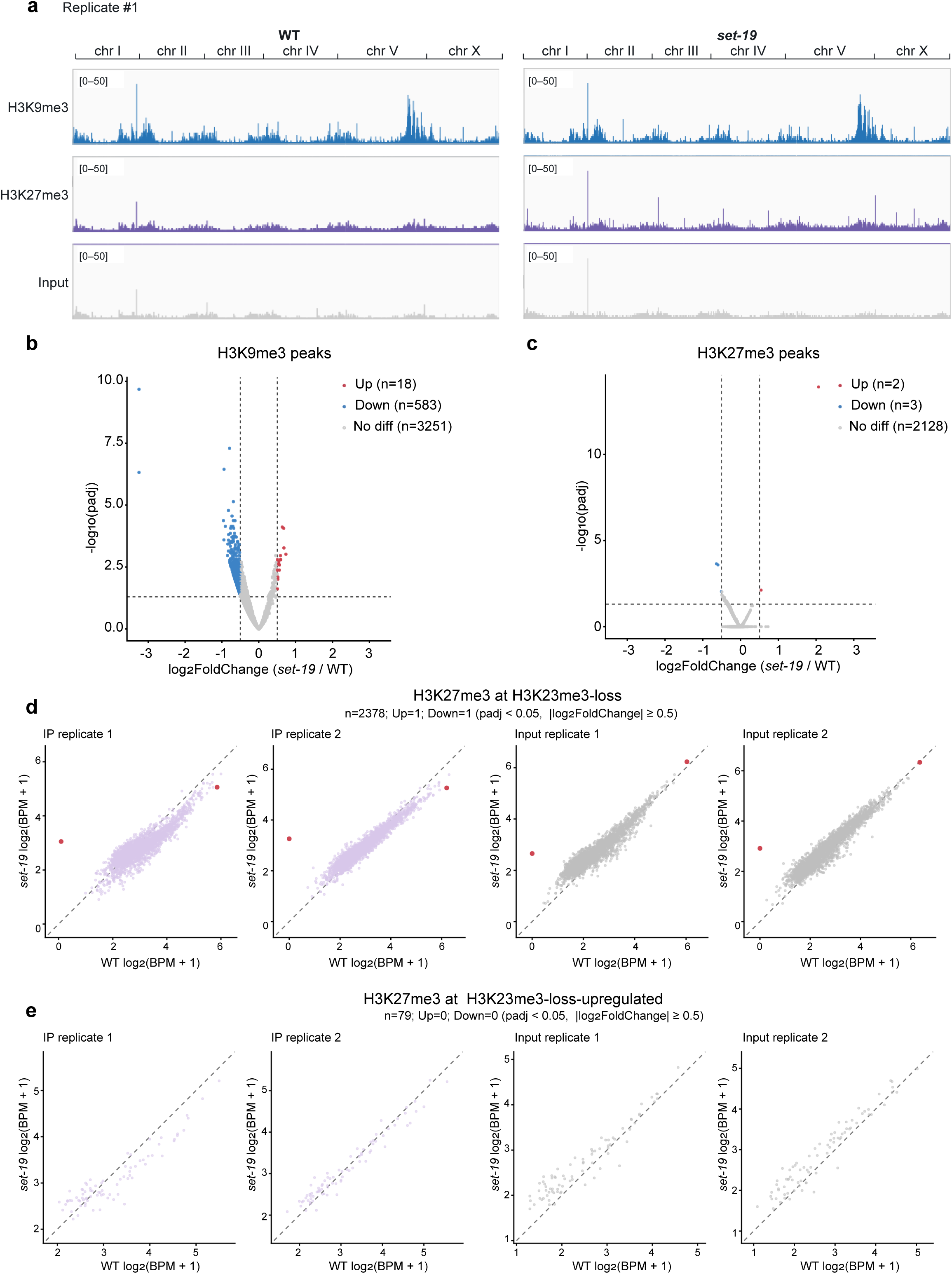
Genome-wide H3K9me3 and H3K27me3 profiles in *set-19* mutants. **a**, Genome browser overview of BPM-normalized H3K9me3 and H3K27me3 ChIP-seq signals and the corresponding Input signals across the six nuclear chromosomes in wild-type (wt) and *set-19(ok1813)* animals. Replicate 1 tracks are shown, using the same visualization range (0–50) for the corresponding wild-type and *set-19* tracks. **b, c,** Differential peak analysis of H3K9me3 (**b**) and H3K27me3 (**c**) between *set-19(ok1813)* and wild-type animals. Volcano plots show log₂FoldChange (*set-19*/wt) versus −log₁₀(adjusted P value [padj]) from DiffBind analysis. Peaks with significantly increased and decreased signals in *set-19* animals are shown in red and blue, respectively (padj < 0.05 and |log₂FoldChange| ≥ 0.5); all other peaks are shown in grey. DiffBind analysis was performed using IP samples only. **d, e,** Scatter plots comparing H3K27me3 IP signals and the corresponding Input signals between wild-type and *set-19(ok1813)* animals at H3K23me3-loss regions (**d**) and at H3K23me3-loss-upregulated regions (**e**). H3K23me3-loss regions were defined by the differential H3K23me3 peak analysis shown in Fig. S5d as peaks with padj < 0.01 and log₂FoldChange < −1 (n = 2,378). The H3K23me3-loss-upregulated regions were defined as H3K23me3-loss peaks overlapping the gene body or strand-specific 2-kb upstream promoter of a gene that was upregulated in *set-19* animals (RNA-seq padj < 0.01 and log₂FoldChange > 1; n = 79). For each region, signals were averaged within a 2-kb window centered on the H3K23me3 peak and are shown as log₂(BPM + 1), with wild-type and *set-19* values plotted on the x- and y-axes, respectively. The two IP replicates and their matched Input replicates are shown separately. Regions with significantly increased or decreased H3K27me3 signals in *set-19* animals, identified by DiffBind analysis of the IP signals (padj < 0.05 and |log₂FoldChange| ≥ 0.5), are shown in red in both the IP and corresponding Input plots.

**Fig. S8.**
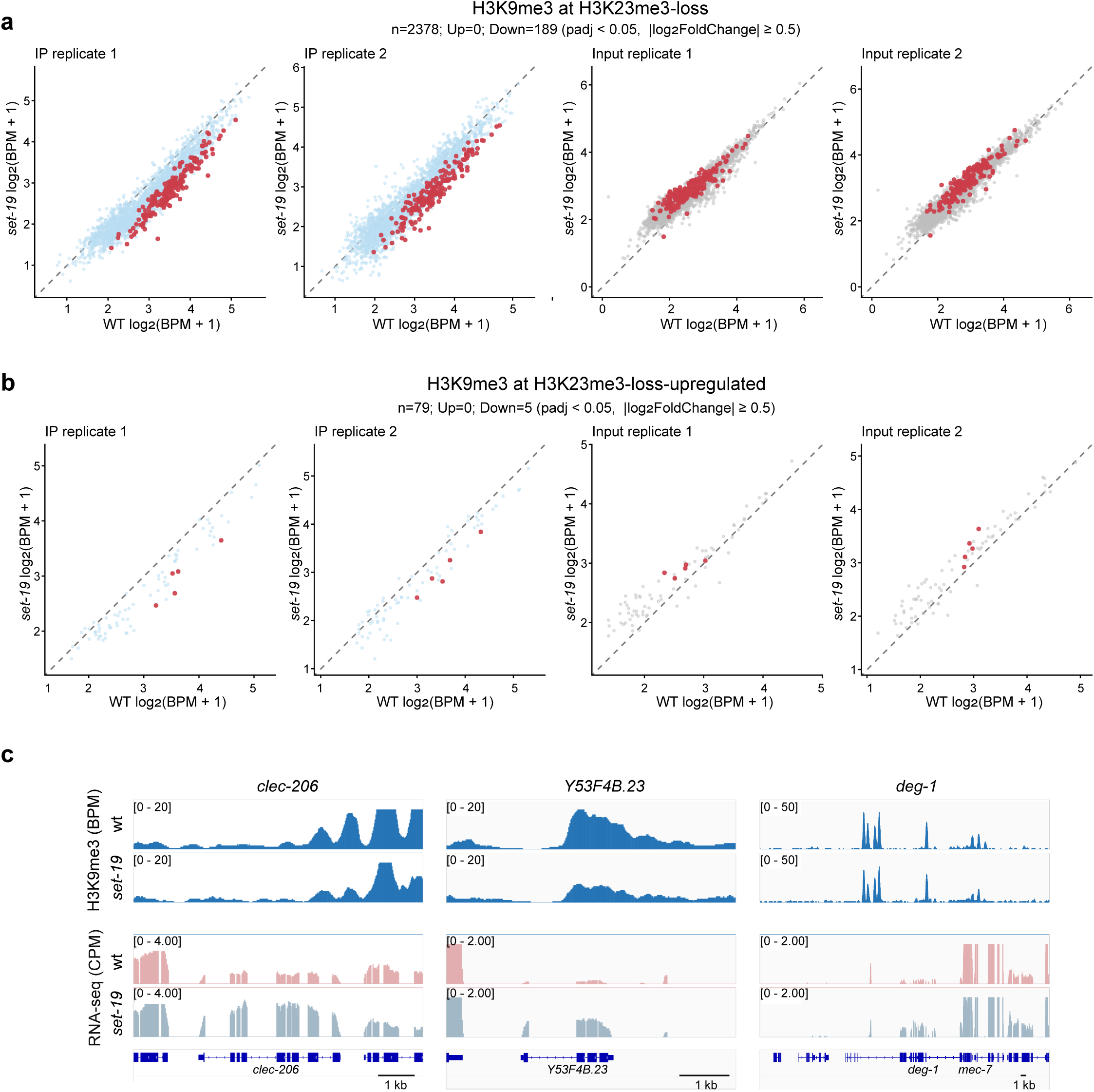
Targeted analysis of H3K9me3 at regions defined by H3K23me3 and transcriptional changes. **a, b**, Scatter plots comparing BPM-normalized H3K9me3 IP and corresponding Input signals between wild-type and *set-19(ok1813)* animals at H3K23me3-loss regions (**a**), and H3K23me3-loss-upregulated regions (**b**). Region definitions and signal calculation and visualization were as described for Fig. S7d, e. Regions with significantly reduced H3K9me3 in *set-19* animals, identified by DiffBind analysis of the IP signals (padj < 0.05 and log₂FoldChange ≤ −0.5), are shown in red in both the IP and corresponding Input plots. **c**, Representative genome browser tracks at *clec-206* and *Y53F4B.23*, which show decreased H3K9me3 signals accompanied by increased RNA-seq signals in *set-19(ok1813)* animals, and at *deg-1*, which shows no detectable H3K9me3 change. H3K9me3 tracks show BPM-normalized IP signals from replicate 1, and RNA-seq tracks show mean CPM-normalized coverage. Signal ranges are indicated in brackets, and gene models are shown below the tracks. Scale bars, 1 kb.

**Fig. S9.**
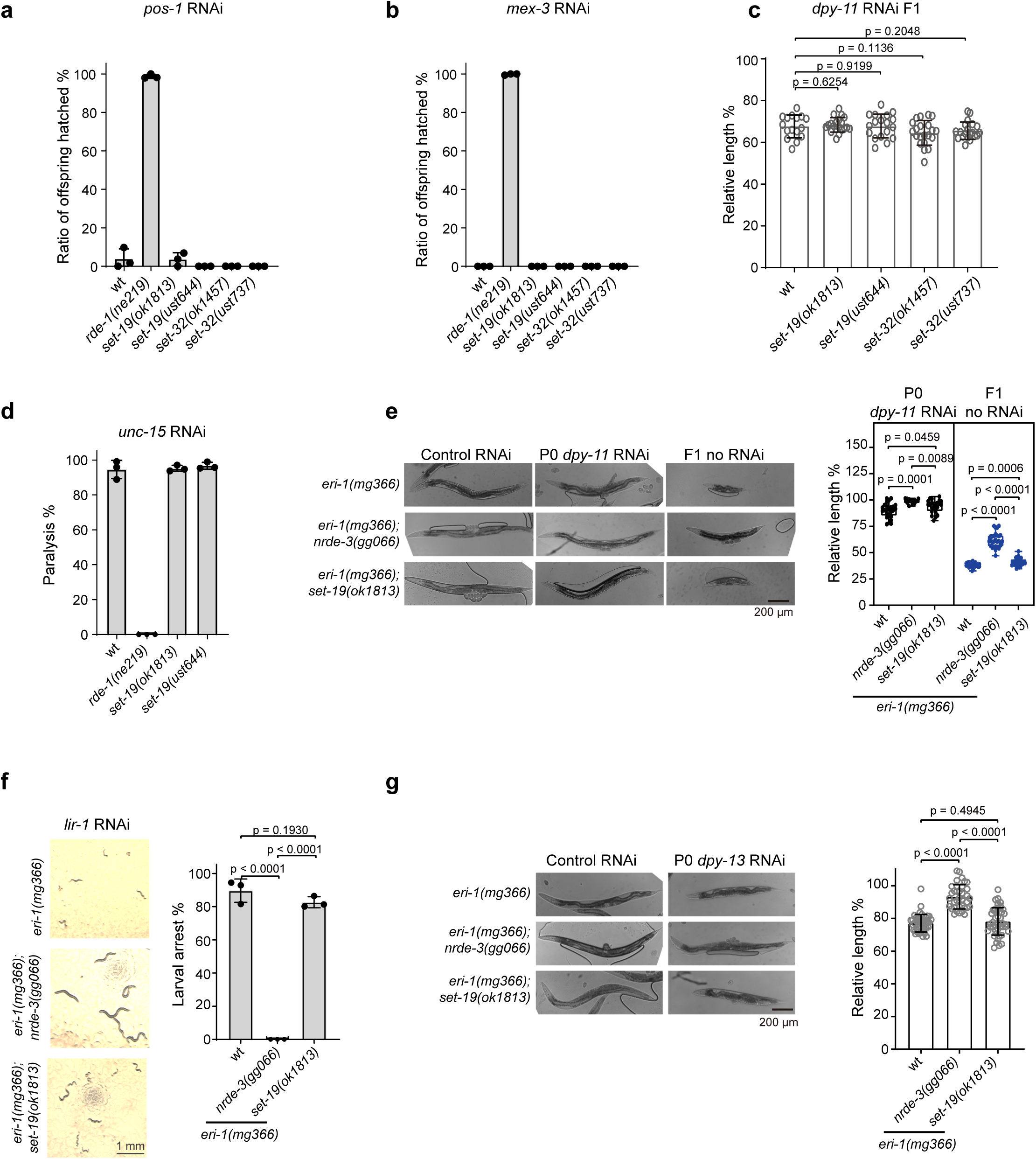
SET-19 is dispensable for feeding RNAi responses. **a, b,** Quantification of embryo hatching rates following feeding RNAi targeting *pos-1* (**a**) or *mex-3* (**b**). Synchronized adult hermaphrodites of the indicated genotypes were exposed to dsRNA-expressing bacteria, and the ratio of hatched offspring was scored. **c,** Analysis of RNAi efficiency using *dpy-11* feeding RNAi. The indicated animals were exposed to *dpy-11* dsRNA until adulthood, and the relative body length of F1 progeny was measured. Individual data points represent single animals. **d,** Quantification of paralysis phenotypes following *unc-15* RNAi in the indicated genotypes. The percentage of paralyzed animals was scored. **e,** Animals of the indicated genotypes were exposed to *dpy-11* RNAi in the P0 generation and transferred to control bacteria. Representative images of P0 and F1 animals are shown (left). The relative body length of P0 and F1 animals was quantified (right). **f,** Larval arrest analysis following *lir-1* RNAi in the *eri-1(mg366)* background. Representative images of larvae are shown (left), and the percentage of animals exhibiting larval arrest was quantified (right). **g,** Analysis of the RNAi response to *dpy-13* feeding RNAi in the *eri-1(mg366)* background. The indicated animals were exposed to dsRNA until adulthood, and the relative body length was measured.

**Fig. S10.**
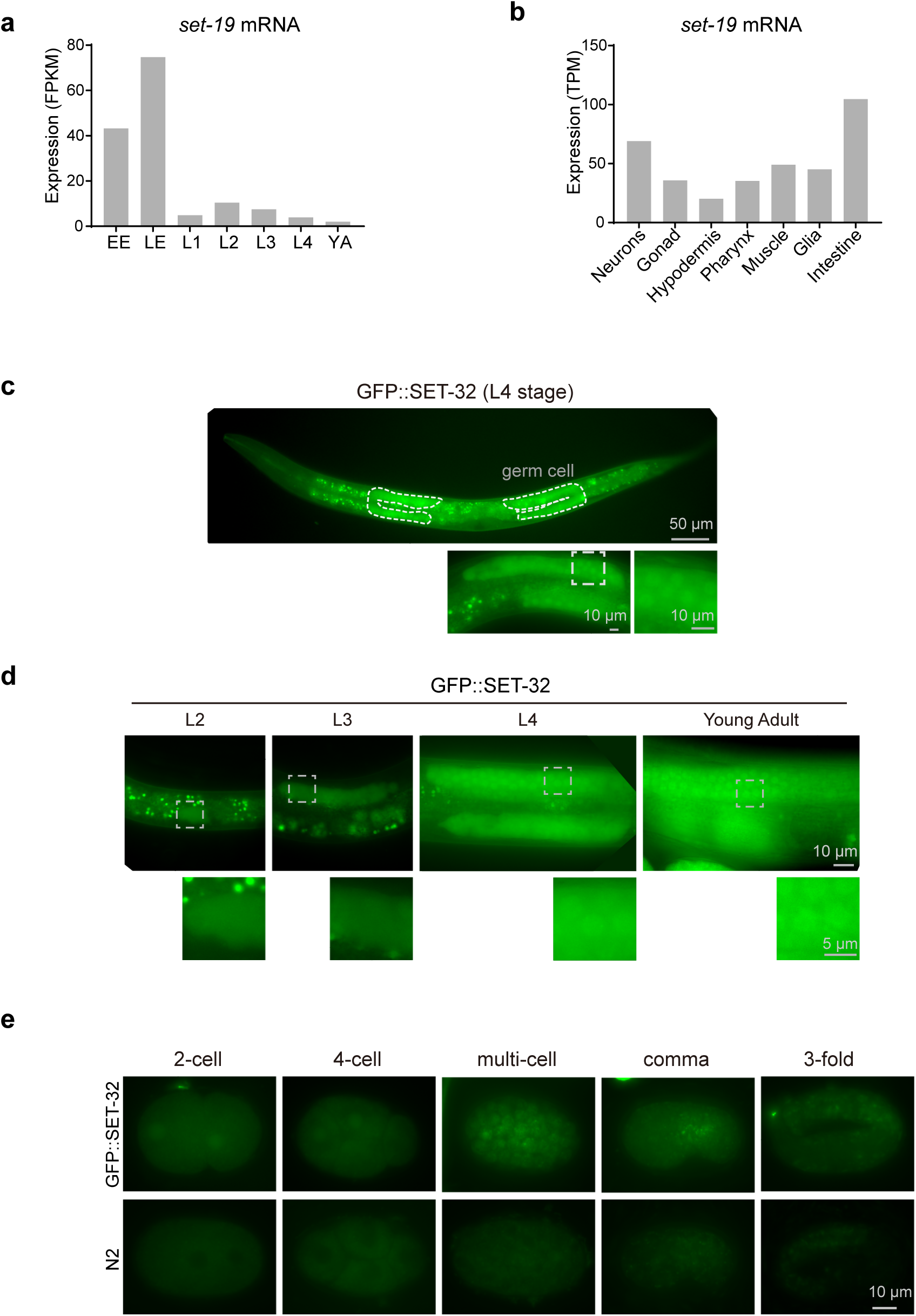
Expression pattern of SET-19 and SET-32. **a,** The expression levels of *set-19* mRNA at different developmental stages. Data were downloaded from WormBase (version WS297). EE, early embryos; LE, late embryos; YA, young adults. **b,** The expression levels of *set-19* mRNA in different tissues at the L2 stage ^67^. **c,** Fluorescence images of L4-stage animals expressing GFP::SET-32. **d,** Subcellular localization of GFP::SET-32 in the germline at different developmental stages. **e,** Subcellular localization of GFP::SET-32 in embryos.

**Table S1.** Strain list used in the work.

| **Strain** | ***Genotype*** |
|---|---|
| N2 | wide type |
| RB1517 | *set-19(ok1813) X* |
| VC967 | *set-32(ok1457) I* |
| yy513 | *pkIs32[pie-1p::gfp::h2b] III* |
| yy528 | *hrde-1(tm1200) III; pkIs32[pie-1p::gfp::h2b] III* |
| OD139 | *ltIs37 [pie-1p::mCherry::his-58 + unc-119(+)] IV* |
| WM27 | *rde-1(ne219) V* |
| GR1373 | *eri-1(mg366) IV* |
| YY182 | *nrde-2(gg091) II;eri-1(mg366) IV* |
| YY189 | *nrde-3(gg066) X;eri-1(mg366) IV* |
| shg695 | *ustSi338[fib-1p::gfp::set-19] II* |
| shg985 | *ustIS205[gfp::set-32] I* |
| shg3155 | *set-19(ust644) X* |
| shg3267 | *set-19(ust703[3xflag::gfp::set-19]) X* |
| shg3270 | *set-19(ust703[3xflag::gfp::set-19]) X; ltIs37 [pie-1p::mCherry::his-58 + unc-119(+)] IV* |
| shg3320 | *set-32(ust737) I* |
| shg3324 | *set-19(ust644) X; ustIS338[fib-1p::gfp::set-19] II* |
| shg3325 | *set-19(ok1813) X; ustIS338[fib-1p::gfp::set-19] II* |
| shg2757 | *set-19(ok1813) X; pkIs32[pie-1p::gfp::h2b] III* |
| shg2758 | *set-32(ok1457) I; pkIs32[pie-1p::gfp::h2b] III* |
| shg2759 | *set-25(n5021) III; pkIs32[pie-1p::gfp::h2b] III* |
| MH1870 | *kuIs54[sur-5:::gfp]* |
| yy520 | *nrde-3(gg066) X; kuIs54 [sur-5:::gfp]* |
| shg483 | *rde-1(ne219) V; kuIs54 [sur-5:::gfp]* |
| shg3522 | *set-19(ust644) X; kuIs54 [sur-5:::gfp]* |
| shg3523 | *set-19(ok1813) X; kuIs54 [sur-5:::gfp]* |
| shg3524 | *set-32(ok1457) I; kuIs54 [sur-5:::gfp]* |
| shg3525 | *set-19(ok1813) X; eri-1(mg366) IV* |

**Table S2.**
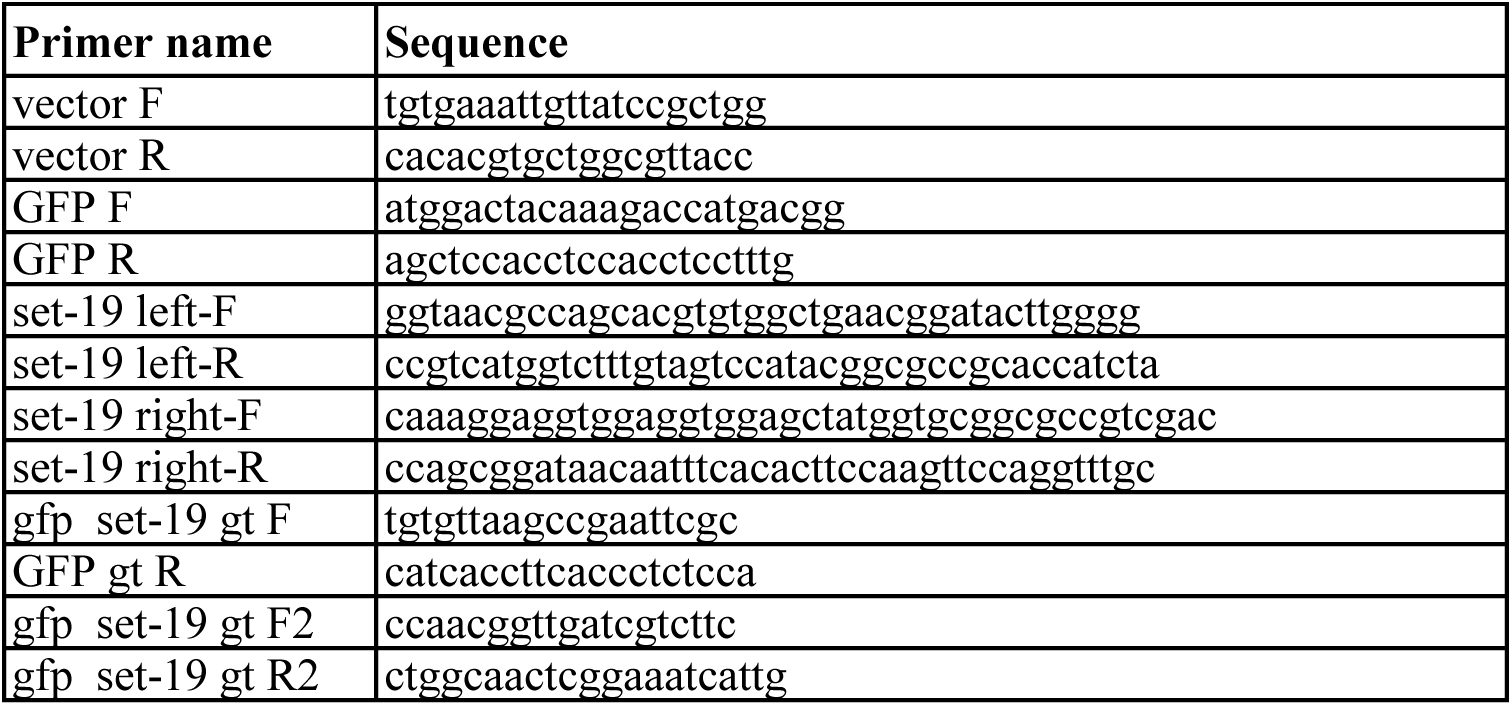
Sequences of primers used in transgenic strain construction.

**Table S3.**
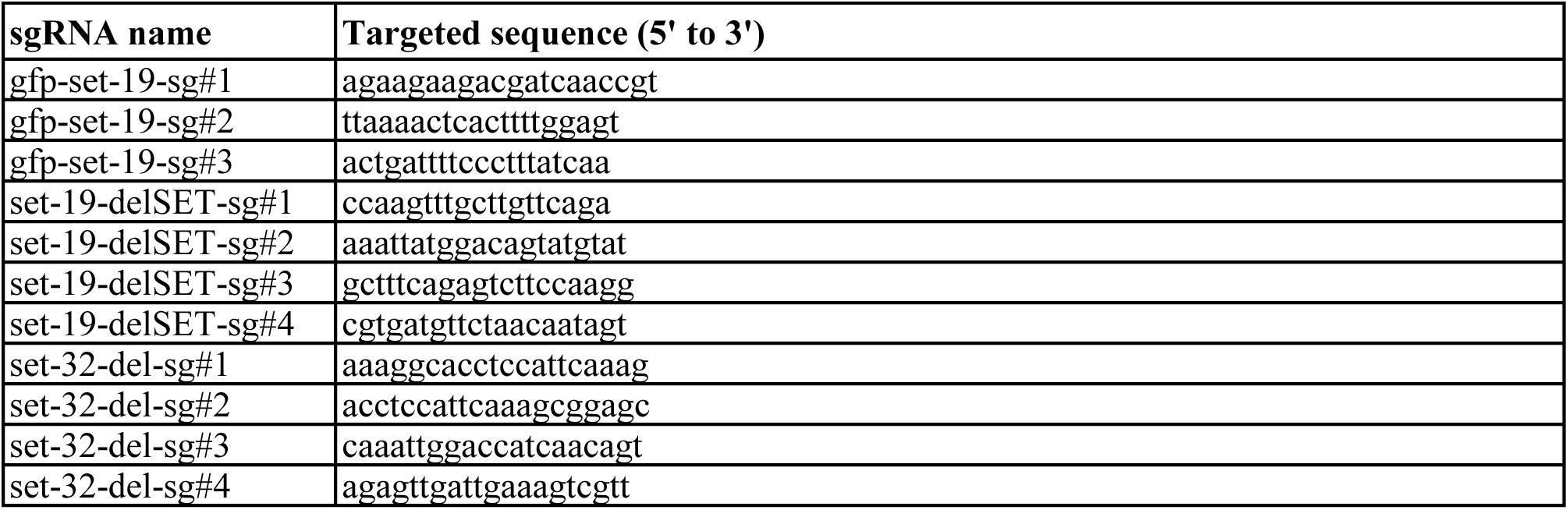
sgRNA targeted sequences for CRISPR/Cas9-directed gene editing technology.

**Table S4.**
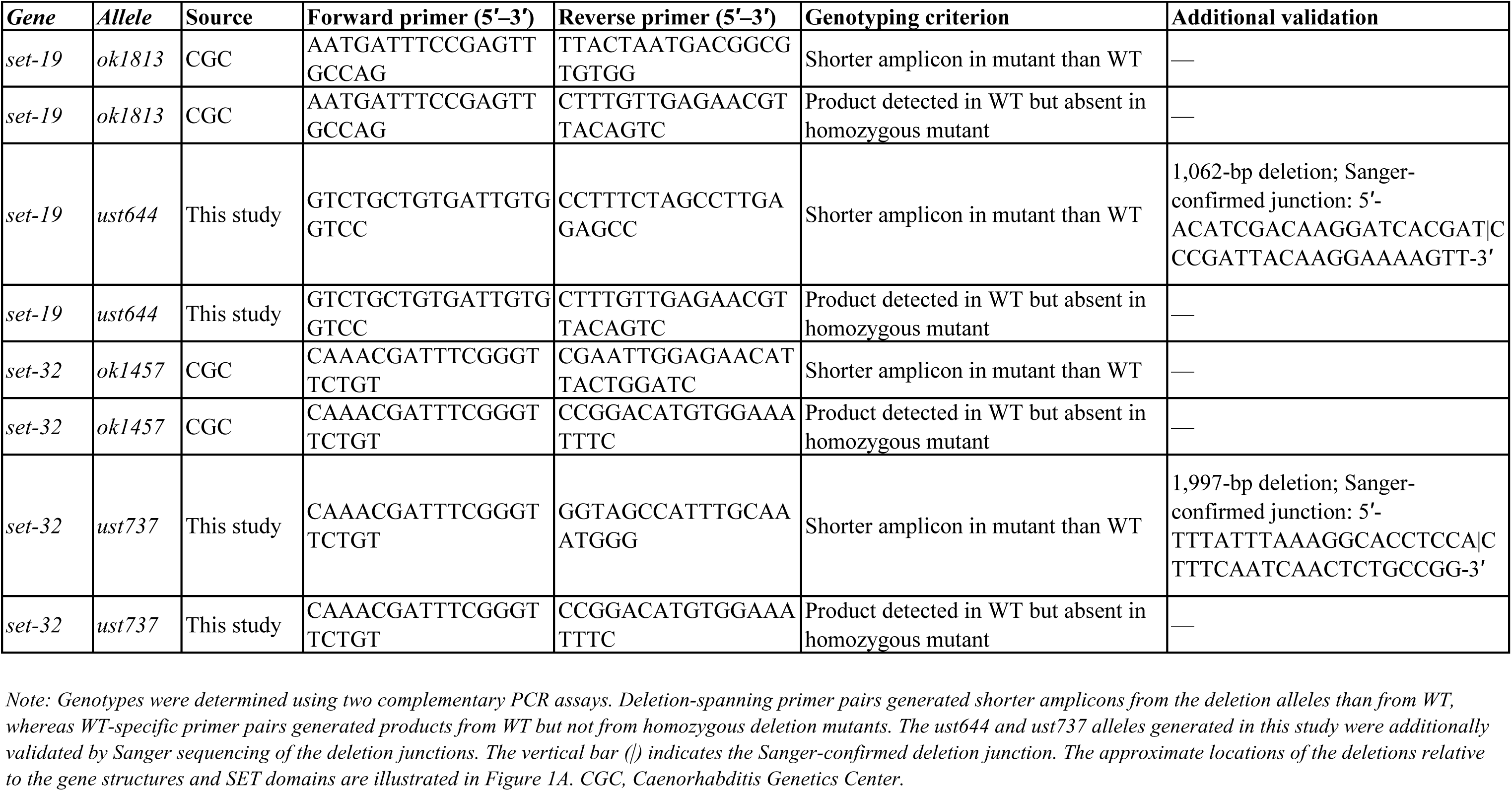
PCR genotyping and validation of the *set-19* and *set-32* deletion alleles used in this study.

**Table S5.** ChIP-seq datasets used in this work.

| **Modifications** | **Antibody** | **Genetic background** | **Stage** | **Resource** |
|---|---|---|---|---|
| H3K4me3 | Abcam, ab8580 | N2 | Adult | SRR7164162 |
| H3K9me3 | Abcam, ab8898 | N2 | Adult | SRR5297105 |
| H3K27me3 | WAKO: 309-95209 | N2 | Adult | SRR7164198 |
| H3K36me3 | Abcam, ab9050 | N2 | Adult | SRR7164186 |
| H3K23me3 | Active Motif, 61499 | N2 | L3-L4 | This work |
| set-19 H3K23me3 | Active Motif, 61499 | *set-19(ok1813)* | L3-L4 | This work |
| H3K9me3 | Active Motif, 39161 | N2 | L3-L4 | This work (Fig. S7, S8) |
| set-19 H3K9me3 | Active Motif, 39161 | *set-19(ok1813)* | L3-L4 | This work (Fig. S7, S8) |
| H3K27me3 | Millipore, 07-449 | N2 | L3-L4 | This work (Fig. S7, S8) |
| set-19 H3K27me3 | Millipore, 07-449 | *set-19(ok1813)* | L3-L4 | This work (Fig. S7, S8) |

